# Robustness of epithelial sealing is an emerging property of local ERK feedbacks driven by cell elimination

**DOI:** 10.1101/2020.03.17.994921

**Authors:** Léo Valon, Anđela Davidović, Florence Levillayer, Mathilde Chouly, Fabiana Cerqueira-Campos, Romain Levayer

## Abstract

While the pathways regulating apoptosis and cell extrusion are rather well described^1,2^, what regulates the precise spatio-temporal distribution of cell elimination in tissues remains largely unknown. This is particularly relevant for epithelia with high rates of cell elimination, a widespread situation during embryogenesis^3–6^ and epithelial homeostasis^7^, where concomitant death of neighbours could impair the maintenance of epithelial sealing. However, the extent to which epithelial tissues can cope with concomitant cell death, and whether any mechanism regulates such occurrence have never been explored so far. Here, using the *Drosophila* pupal notum (a single layer epithelium) and a new optogenetic tool to trigger caspase activation and cell extrusion, we first show that concomitant death of clusters of at least three cells is sufficient to transiently impair epithelial sealing. Such clustered extrusion was almost never observed *in vivo*, suggesting the existence of a mechanism preventing concomitant elimination of neighbours. Statistical analysis and simulations of cell death distribution in the notum highlighted a transient and local protective phase occurring near every dying cell. This protection is driven by a transient activation of ERK in the direct neighbours of extruding cells which reverts caspase activation and prevents elimination of cells in clusters. Altogether, this study demonstrates that the distribution of cell elimination in epithelia is an emerging property of transient and local feedbacks through ERK activation which is required to maintain epithelial sealing in conditions of high rate of cell elimination.

---

Epithelial cell elimination is driven by extrusion, a succession of remodeling steps removing one cell from the epithelial layer while maintaining epithelial sealing^1,8^. We first asked whether concomitant extrusion of several neighbours impairs the maintenance of epithelial sealing. As such, we developed a UAS-optoDronc *Drosophila* line, which can trigger rapid caspase activation through blue light-induced clustering of Caspase9 (**Figure 1a, Figure S1a**). Expression of optoDronc in fly eyes (GMR-gal4 eye-specific driver) is sufficient to trigger cell death upon blue light exposure and can be rescued by expression of the downstream effector caspase inhibitor p35 (**Figure S1b**). We then used the *Drosophila* pupal notum to assess the efficiency of the construct in triggering epithelial cell elimination. Blue light exposure of clones expressing optoDronc triggers elimination of the majority of cells in less than one hour (**Figure 1b, Figure S1c,e, movie S1**). OptoDronc-triggered extrusions are similar to physiological extrusions in the pupal notum, albeit slightly faster (**Figure S1f**), and require effector caspase activation (**Figure S1d,e, movie S1**) like physiological extrusions in the notum^9,10^. We then induced concomitant elimination of group of cells of various sizes and shapes by expressing optoDronc in clones. While extrusions of single cells or several cells in lines occurred normally (**Figure 1b,e,f, movie S2 left**), concomitant extrusion of three cells or more in cluster led to aberrant extrusion: cells initiate contraction then relax transiently (**Figure 1c-f, movie S2 right**) and eventually close the gap through a process akin to wound healing (**Figure 1c**, E-cad accumulation at vertices, **movie S2 right**). Aberrant extrusions correlated with transient flow of injected extracellular fluorescent Dextran in between cells at the level of adherens junctions (**Figure 1g,i,j, movie S3 bottom**), suggesting that epithelial sealing is transiently impaired. Dextran flow however was not observed for cells eliminated in lines (**Figure 1g,h,j, movie S3 top**). Altogether, we conclude that concomitant extrusion (<30 min) of three cells or more in cluster leads to transient loss of epithelial sealing and is thus detrimental for the tissue.

**Figure 1:**
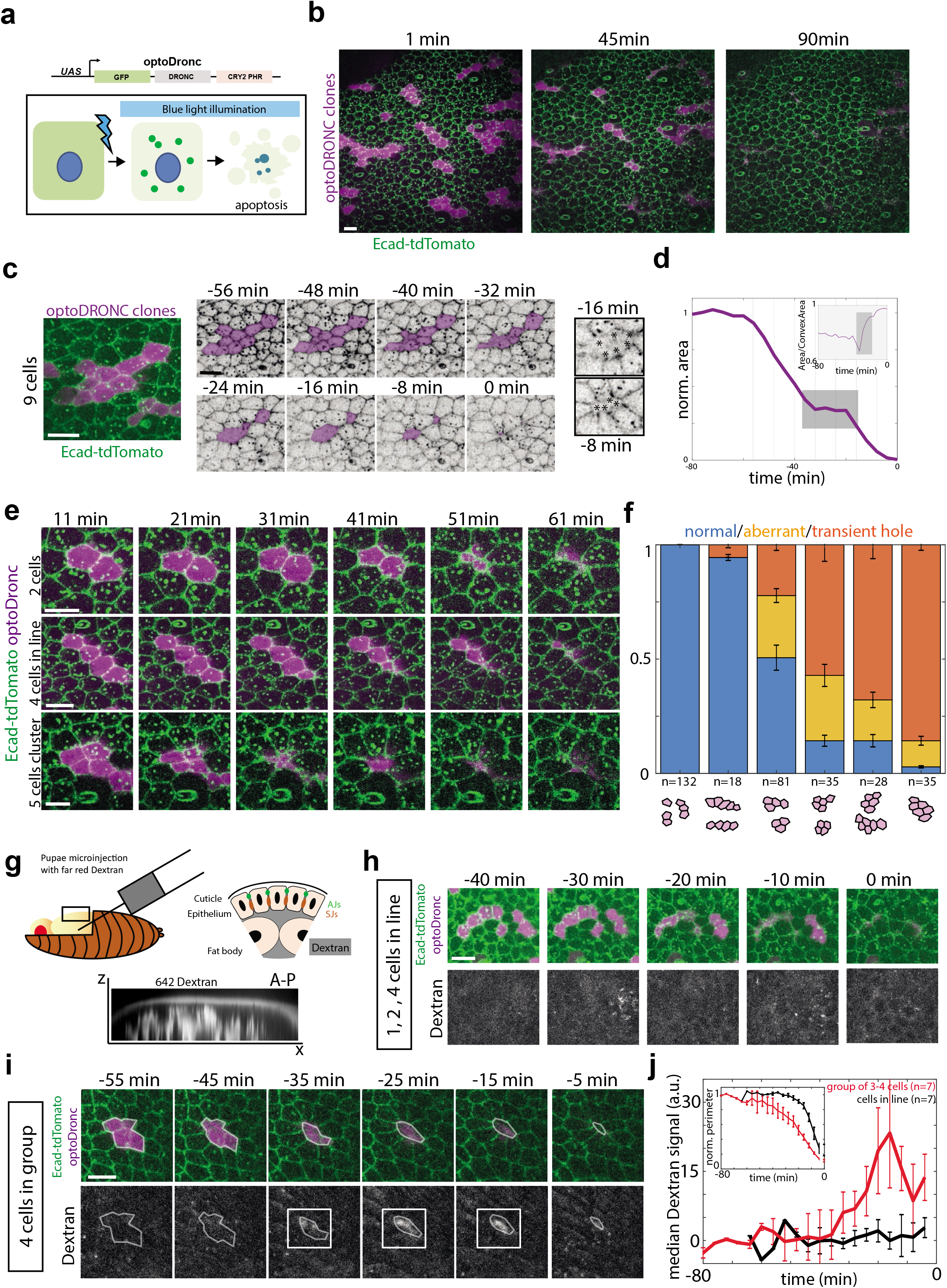
Extrusion of a cluster of more than three cells is sufficient to impair epithelial sealing. **a:** Schematic of the UAS-optoDronc construct. The cDNA of Dronc (*Drosophila* Caspase9) is fused to eGFP in N-ter and the blue light sensitive protein CRY2-PHR in C-ter. Upon blue light exposure, optoDronc clusterises (green dots) and triggers caspase activation and apoptosis. **b:** Snapshots of a live pupal notum expressing optoDronc in clones (magenta, local z-projection) and E-cad-tdTomato. Most of the clones disappear after 60 minutes of blue light exposure. Scale bar= 10μm. **c:** Elimination of an optoDronc clone of nine cells. Snapshots of inverted E-cad-tdTomato signal. Time 0 is the termination of clone elimination. Clone contour rounds up and relaxes at −32 min and is followed by wound healing. Inset on the right shows E-cad accumulation at tricellular junctions during wound healing. Scale bar= 10μm. **d:** Evolution of the clone area shown in **c**. The grey zone corresponds to the relaxation phase and is followed by wound healing. The lines corresponds to the timing of the images shown in **c**. Inset shows increased of clone solidity (area/convex area) during the relaxation and wound healing phases (grey zone). **e:** Snapshots of clones of different sizes expressing optoDronc upon blue light exposure. Note the transient relaxation at 41 min for 5 cells cluster. Time 0 is the onset of blue light exposure. Scale bars= 10μm. **f:** Quantification of the proportion of normal and aberrant extrusions (extrusions followed by transient relaxation or E-cad accumulation at vertices) and/or transient hole (large relaxation and E-cad accumulation at vertices) observed for clone disappearance of different sizes and topologies (see schematic below). n=number of clones. Error bars are 95% confidence interval. **g:** Injection of far red Dextran 10,000 MW in pupal notum. Bottom shows a transverse view of Dextran signal in the pupae (x-antero-posterior, Z-apical-basal). Schematic showing the localisation of Dextran according to the epithelium (AJs: adherens junctions, SJs: septate junctions). Note that septate junctions are located basally to adherens junctions. **h:** Local projections of optoDronc clones composed of one to several cells in lines after Dextran injection (bottom). Time 0 is the end of clone elimination. No dextran appears at the level of adherens junctions during clone elimination. Scale bar= 10μm. **i:** Local projections of an optoDronc clone composed of four cells in cluster after Dextran injection (bottom). Time 0 is the end of clone elimination. During the relaxation phase (−35min) dextran appears at the level of adherens junctions (white squares). Scale bar= 10μm. **j:** Quantification of the far red Dextran signal during optoDronc clone elimination (in line, black, or in clusters of 3-4 cells, red). The curves are the median +/- s.e.m.. Top inset shows the average clone area during their elimination (note that the elimination is overall slower for clusters, red curve).

Given the rate of cell elimination and assuming that cell eliminations are independent (Poisson process), concomitant elimination of 3 cells or more in cluster should occur several times per movie (See **Methods** for details, more than 2 events per movie of 20 hours). Yet, we very rarely observed such clusters during notum morphogenesis (<1 case per movie, **Figure 4k**, n=5 pupae). We therefore checked whether cell death distribution was indeed following locally a Poisson process in the pupal notum. We focused on the posterior region of the notum where cell death distribution is rather uniform in time and space (**Figure 2A, movie S4**) to neglect as much as possible the impact of tissue patterning on the rate of cell elimination. We first characterised the distribution of cell death by calculating for every cell elimination the local density of cell death at different distances and different times from the eliminated cell (**Figure 2b**). We obtained a map of local death density for every movie (**Figure 2c**) which was then compared to 200 simulations of the death distribution assuming a Poisson process at the same rate of cell elimination. The difference between the simulated map and experimental map was then used to check local differences in the distribution (**Figure 2c, Figure S2a**) and averaged for 5 nota (**Figure 2d**). Strikingly, there was in our experimental data a significant reduction in the density of cell elimination in the vicinity of each dying cell (<7μm) in a short time window (starting at 10 minutes and up to 60 minutes) compared to the simulated distributions. This suggests that cell death distribution in the posterior region of the pupal notum does not strictly follow a Poisson process, and that a transient (~one hour) and local (~one cell diameter) refractory phase appears in the vicinity of each dying cell. To confirm this bias, we then used a closest neighbour analysis (**Figure 2e**): for each cell elimination we detected the nearest elimination in different time windows. While the distribution in late time windows overlapped (1h20’ to 2h20’, 2h20’ to 3h20’ after cell death **Figure 2 f,g**), there was a significant decrease of the probability of cell death for the first 60 minutes (20 to 80 minutes) following each cell elimination at a distance of 0-10 μm (~one cell diameter, two folds reduction, p=0.008 for distances<5μm). Finally, to confirm the existence of a local dispersion of cell death, we performed a classical p-value tests for dispersion using K-functions (see **Methods** and ^11^). This method detects potential temporal and spatial dispersion in a distribution. To validate the approach, we first compared the distribution of one of our experiments with the one of a simulated Poisson process with the same effective death intensity. We also compared the experimental distribution with the one of a simulated process with same intrinsic death intensity (see **Methods**) adding a local and transient inhibition of cell death near every dying cell (5 μm range, 40 min inhibition starting 10 min after the initial death, see **Methods**). While no significant dispersion appears in the Poisson distribution, a significant dispersion peak appears both for the experimental distribution and the simulated one with local inhibition (**Figure 2h**, simulated maps are averaged of 20 simulations), and similar dispersion peaks were found for the other movies analysed (**Figure S2b, Figure 2i**). Altogether, we concluded that cell elimination is followed by a transient and local refractory phase (~one cell distance, with a delay of ~10-20 minutes and lasting ~60 minutes) that reduces locally and transiently the probability of cell death (up to 2 folds).

**Figure 2:**
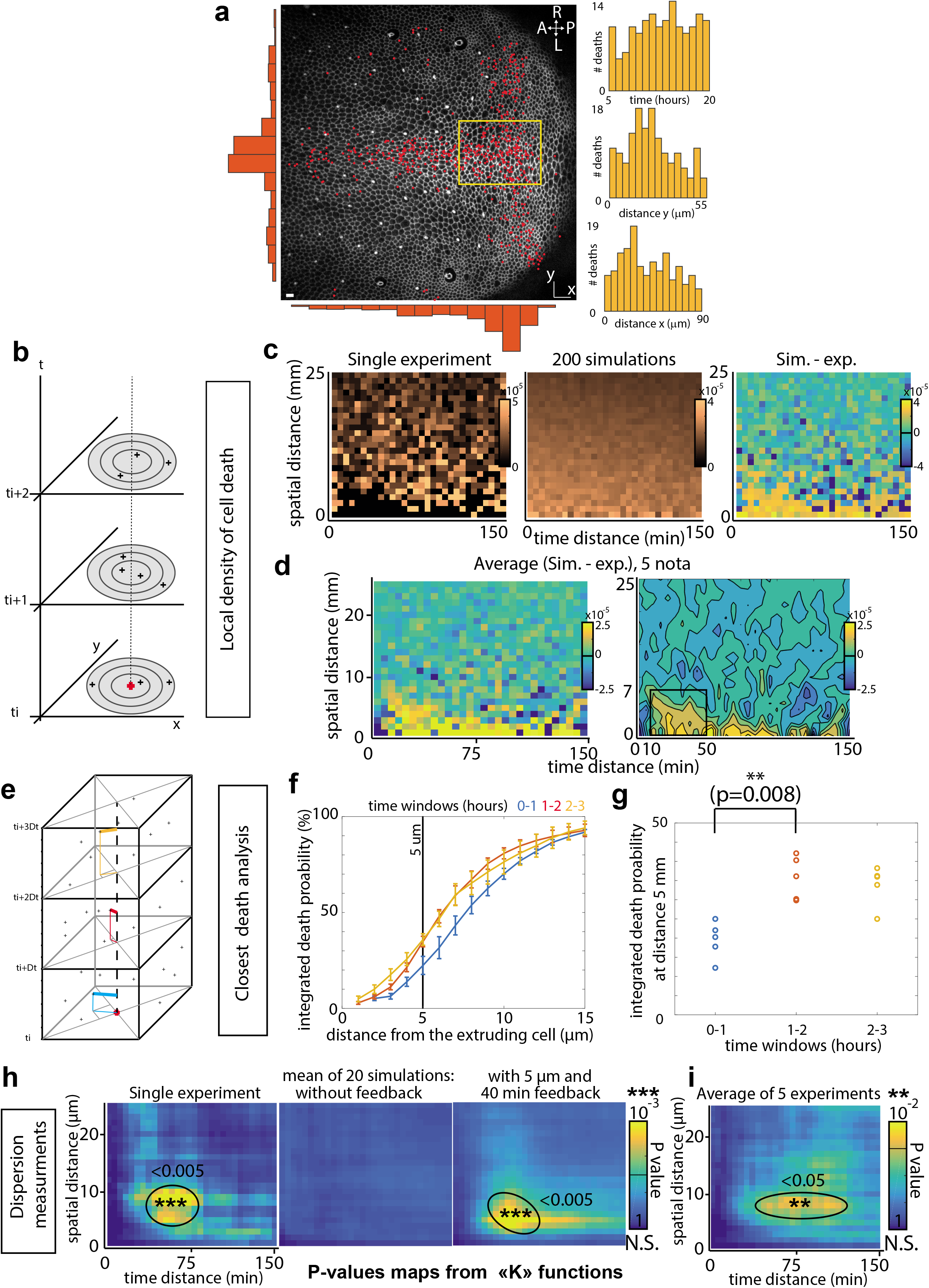
There is a transient and local refractory phase for cell elimination following each cell death. **a:** Snapshot of a local projection of a pupal notum expressing E-cad-GFP. Red dots are showing every cell extrusion occurring over 21 hours. A: Anterior, P: Posterior, L: Left, R: Right. Scale bar= 10μm. Orange histograms show the spatial distribution of cell death over left-right axis (left) or along AP axis (bottom). Right histograms show the temporal distribution of cell death (top) and its spatial distribution (middle and bottom) in the yellow rectangle region (used for further analysis). **b:** Scheme explaining the measurement of spatial and temporal distance between death events. For each death event, the density of cell death in a disc (number of death events divided by disc surface) at a given spatial distance is calculated for each time point. **c:** Local cell death density at different spatial (y axis) and temporal distances (x axis) from a dying cell for one movie. The middle map shows the average map obtained for 200 simulations of a Poisson process with the same cell death intensity. The right map shows the difference between the simulated and the experimental distributions (“yellow island” shows that death at short distances are under-represented in the experiment compared to the simulations). **d:** Average of the difference between experimental maps and the corresponding simulated maps (simulation minus experimental distributions, see **Figure S2** for details, 5 movies). The right map is obtained after median filtering. Note the bottom left yellow domain with lower expected number of cell deaths compared to simulations (black square, 7μm~one cell diameter, and from ~ 10 to 60min). **e:** Analysis of cell death distribution through a closest neighbour approach. Time is subdivided in arbitrary windows (here 1 hour, starting from t=20 minutes) and for each time window the spatially closest cell death is found. **f:** Cumulative plots of the probability to find the closest death at a given distance for different time windows (20min to 1hour 20, 1h20 to 2h20, 2h20 to 3h20). The curves are averages of 5 movies, error bars are s.e.m.. Note that only the blue curve detaches from the others, representing what happens 20 to 80 minutes after cell death. **g:** Details of the values obtained for a distance of 5μm (~ 1 cell distance) for each time window (one dot = one movie). There is a two-folds reduction of the probability to have cell death occurring during the first hour after cell elimination for distances from 0 to 5 μm to the dying cell. **h:** Analysis of dispersion in the death distribution using maps of dispersion p-value calculated with K-functions (see **Methods**). Y-axis: spatial distance between death events, X-axis: time delay between death events. Pseudo colour is the p-value (yellow=significant dispersion, blue=no significant dispersion). The left map corresponds to the experimental distribution used in **2c**, middle map is the mean of the maps from 20 simulations of a Poisson process with the same death intensity; the right map is the mean of the maps of 20 simulations of a random process including a cell death refractory phase following each cell death starting with a delay of 10min and lasting 40min at a distance of 5μm (see **Methods**). **i:** Averaged p-value map for 5 WT movies (see **Figure S2** for details).

These results suggest the existence of active mechanisms preventing the elimination of the neighbours and generating this refractory phase. Effector caspase activation systematically precedes and is required for every cell extrusion in the pupal notum^9,10^. We therefore tracked caspase dynamics using a GFP live sensor of effector caspase activity (GC3Ai, ^12,13^) and used the rate of GFP signal accumulation as a proxy for caspase activity (see **Methods**). We frequently observed neighbours simultaneously activating caspases (**Figure 3a, movie S5**), however extrusion of the first cell led to reversion of caspase activity in the neighbours which then remained in the tissue (**Figure 3a,b,c**, caspase activity goes back to basal levels in the neighbours). This suggested that cell extrusion could inhibit caspase activity and elimination of the direct cell neighbours. We next asked which mechanism could modulate the dynamics of caspase activation and bias the distribution of cell elimination. Recently we showed that the EGFR/ERK pathway is a central regulator of cell elimination and caspase activity in the pupal notum^10^, mostly through the inhibition of the pro-apoptotic gene *hid*^10,14,15^. Moreover, EGFR/ERK can be activated by tissue stretching and downregulated by tissue compaction^10^. Using a live sensor of ERK (miniCic-mScarlet, adapted from^10^, nuclear accumulation of mScarlet at low ERK activity and nuclear exclusion at high ERK activity, **Figure 3d**), we monitored ERK dynamics near extruding cells. Strikingly, we observed systematically a transient activation of ERK in the direct neighbours of every dying cell (**Figure 3d,f, movie S6, Figure S3a,b, movie S7**, activation also observed with EKAR, a ERK FRET sensor^16^). This activation was concomitant with the transient stretching of the neighbouring cells triggered by cell extrusion (**Figure 3 e,f**), lasted for ~ 60min and was restricted to the direct neighbours of the dying cell (**Figure 3f**). ERK activation does not correlate with pulses of Calcium (**Figure S4a,b, movie S8**), does not require active secretion of EGF/Spitz from the dying cells (**Figure S4c,d, movie S9**, contrary to enterocytes elimination in the fly midgut^17^) and can be mimicked by laser induced cell elimination (**Figure S4e,f, movie S10**). We previously showed that stretch-induced survival in the notum required EGFR^10^. Accordingly, ERK activation near dying cells is completely abolished upon EGFR depletion in the tissue (**Figure 3g,h, movie S11**, RNAi previously validated^10^). Moreover, EGFR is only required in the neighbouring cells and not in the dying cell (**Figure S3e**). Altogether, this suggested that ERK activation could be driven by the stretching induced by cell extrusion through EGFR activation, although at this stage we cannot exclude other contact dependent mechanisms. Importantly, these pulses of ERK are not restricted to the pupal notum as similar dynamics were observed near dying larval accessory cells in the pupal abdomen (**Figure S3 c,d, movie S12**).

**Figure 3:**
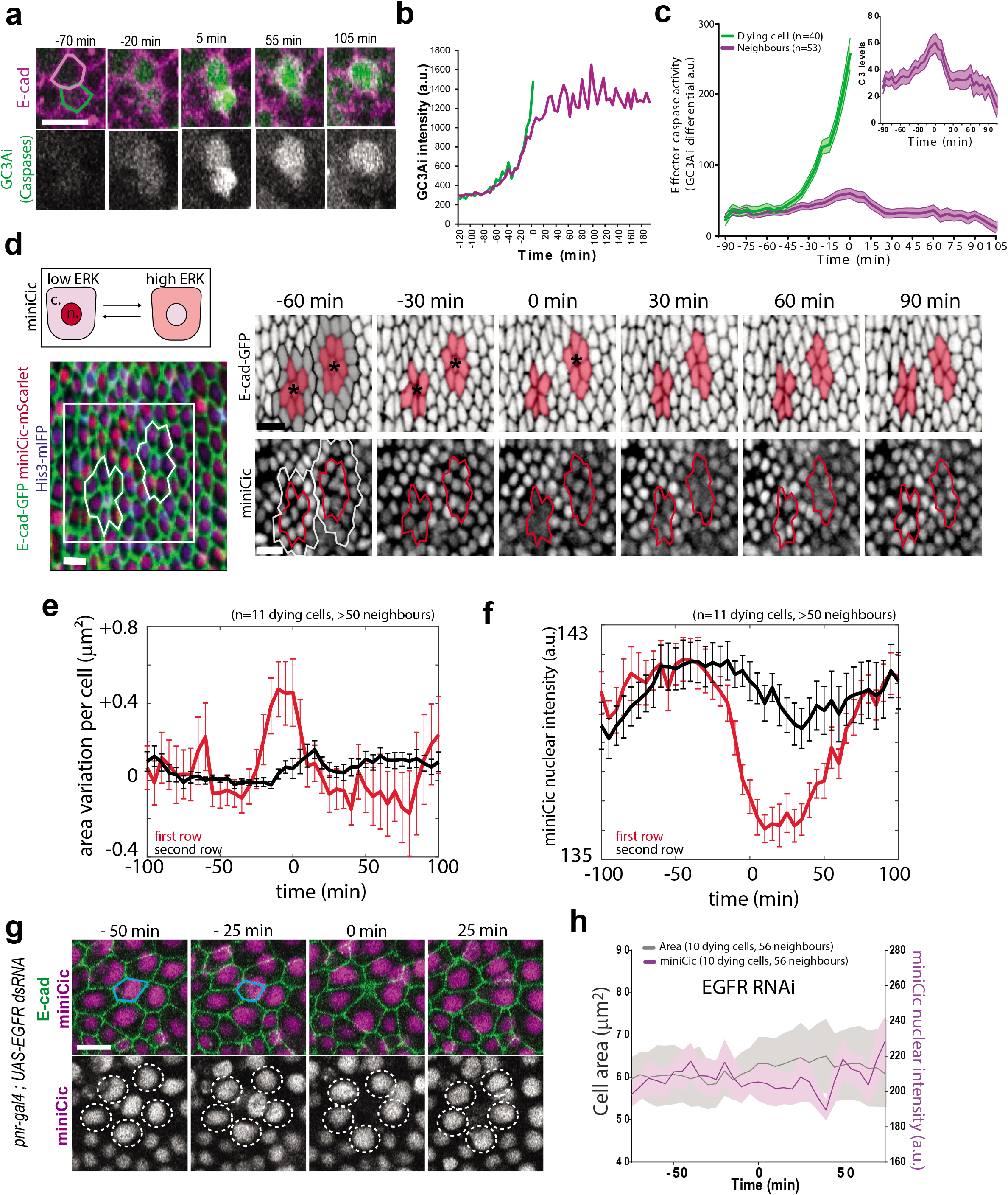
Transient ERK activation in cells neighbouring an extruding cell. **a:** Snapshots of local projections of a pair of cells in the pupal notum tagged with E-cad-tdTomato and expressing the effector caspase sensor GC3Ai (green). Time 0 is the termination of extrusion of the bottom cell (green contour). Scale bar=10μm. **b:** Intensity of GC3Ai in the eliminated cell (green) and its neighbour (purple). Note that GC3Ai signal plateaus in the neigbour after the first cell is eliminated. **c:** Averaged GC3Ai differential (a proxy of effector caspase activity, see **Methods**) for dying cells (green) and its neighbours (purple). Neighbours undergo transient caspase activation followed by reversion. Time 0 is the termination of the first cell extrusion. Light coloured areas show s.e.m. The top inset shows the purple curve with higher magnification. **d:** top: schematic showing the localisation of miniCic (n.: nucleus, c.: cytoplasm) upon modulation of ERK activity. Bottom: Snapshot of the posterior region of the pupal notum with two dying cells (red: miniCic-mScarlet, green: E-cad-GFP, blue: His3-mIFP). Right: snapshots of two dying cells (black stars) showing E-cad signal (inverted colour, top) and miniCic (bottom). Orange zones (red line for miniCic) are the first raw of neighbouring cells, grey zones (white line for miniCic) mark the second raw. Scale bars=10μm. **e:** Averaged cell apical area variation in the first row (red) and second row (black) of cells neighbouring an extruding cell. Time 0 is the termination of extrusion. Error bars are s.e.m.. **f:** Averaged nuclear miniCic intensity in the first row (red) and second row of cells (black). Error bars are s.e.m.. **g:** Snapshots of a *pnr-gal4; UAS-EGFRdsRNA* pupae (local projections) expressing E-cad-GFP, miniCic-mScarlet and His3-mIFP (not shown). The blue line shows a dying cell. The white doted circles show miniCic signal in the nuclei of the neighbouring cells. Scale bar=10μm. **h:** Averaged nuclear miniCic intensity (purple) and cell apical area (black) in EGFR depleted nota in the cells neighbouring an extruding cell (Time 0 is the termination of extrusion). Light areas are s.e.m..

We then asked whether ERK pulses were indeed required for caspase reversion in the neighbours. We first correlated ERK dynamics with caspase activity. We observed a significant positive correlation between ERK activity and caspase inhibition with an averaged lag-time of 15 minutes (**Figure 4a-c, movie S13** cross-correlation between nuclear miniCic and GC3Ai differential, r^2^=0.37). Moreover, caspase reversion in the neighbouring cells was abolished upon EGFR depletion (**Figure 4d-f, movie S14**). Thus, ERK pulses precede and are required for caspase inhibition in the neighbouring cells following each cell death. The localisation and duration of ERK activation (1 row of cells, ~one hour, **Figure 3f**) and the delay with caspases inhibition (15 minutes) were remarkably similar to the death refractory zone we observed near every cell elimination (<7μm, 10-20 minutes delay, duration of 60 minutes, **Figure 2**). We therefore checked whether ERK pulses could modulate the spatio-temporal distribution of cell elimination in the posterior region of the notum. Strikingly the refractory phase we found in WT pupae (**Figure 2**) was not observed in EGFR depleted nota either using the local density of cell death compared to simulations (**Figure 4g,h, movie S15**), using the closest neighbours analysis (**Figure 4i**) or using the p-value dispersion test (no significant dispersion peak, **Figure 4j, Figure S2d**). This suggested that EGFR and ERK pulses are indeed required to generate the transient and local death refractory phase. In absence of local inhibition,we expected to observe spontaneous occurrence of clusters of cell elimination. Accordingly, we observed frequent concomitant extrusion of clusters of cells upon EGFR depletion in the notum (**Figure 4k, movie S16**, >3 cells, ~4 events per movie, n=9 pupae). These clusters undergo abortive extrusion, transient relaxation followed by wound healing (**Figure 4m, movie S16**, E-cad accumulation at vertices), similar to the abortive extrusions triggered by optoDronc cell clusters (**Figure 1c-f, movie S2**). Altogether, this showed that ERK pulses are required to reverse caspase activation near dying cells and to prevent elimination of cells in clusters. This feedback is essential to maintain epithelial sealing in conditions with a high rate of cell elimination (**Figure 4m**).

**Figure 4:**
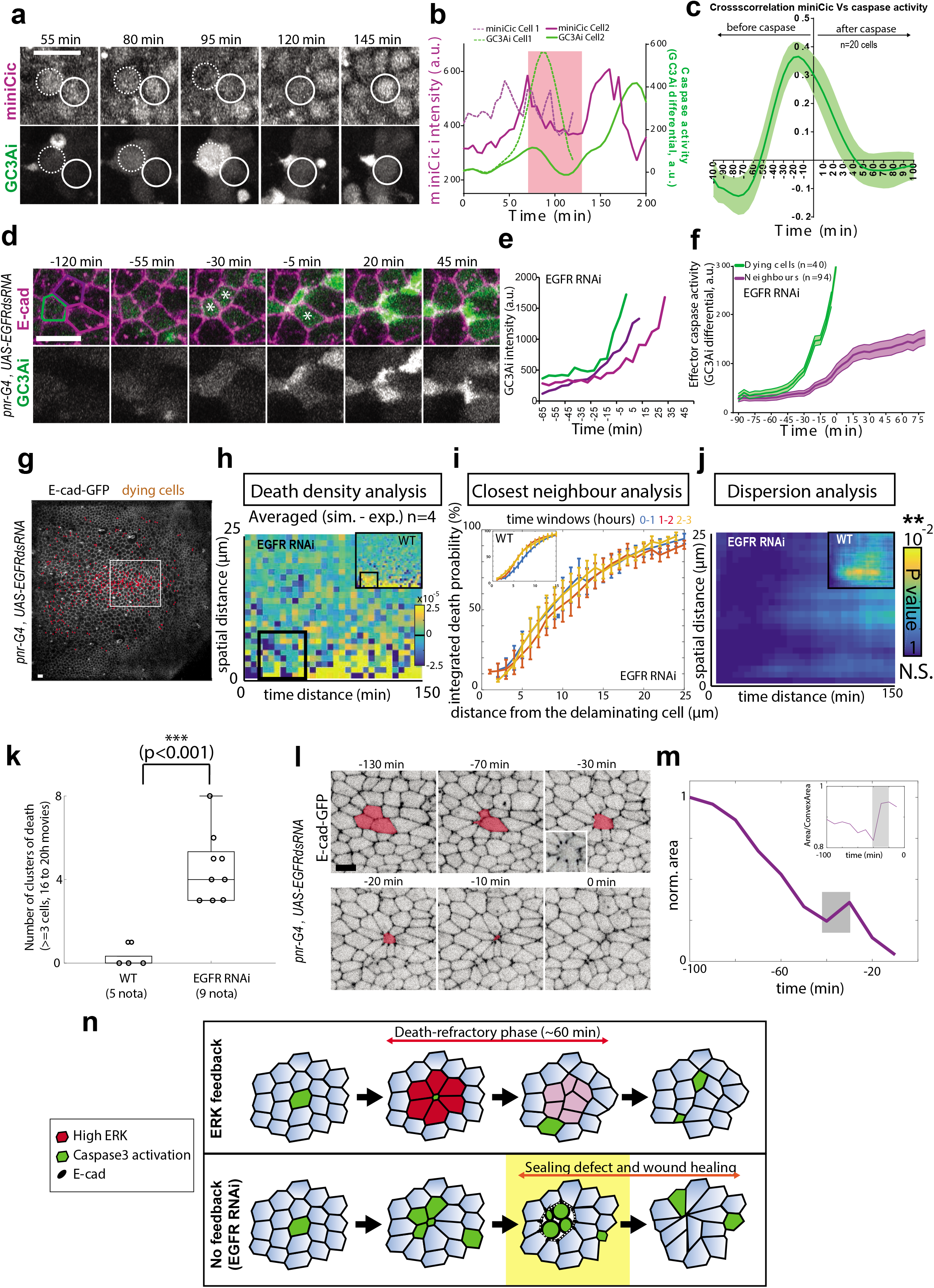
ERK pulses are required for caspase reversion and to prevent clusters of cell elimination. **a:** Snapshots of local projections of a pupal notum expressing miniCic and GC3Ai. White dotted circles show the first dying cell, white circles show one of its neighours dying later. Time is in agreement with the curves shown in **b. b:** Nuclear miniCic signal (magenta) and GC3Ai differential (green, caspase activity) in the first dying cell (dotted lines) and its neighbours (plain lines) shown in **a**. The red region highlights the pulse of ERK (decrease of miniCic) in the neighbours and the subsequent transient reversion of caspase activity. **c:** Averaged normalised cross-correlation between miniCic nuclear signal and caspase activity (GC3Ai differential). Peak at −15 min indicates a 15 minutes delay between ERK activation and caspase reversion. **d:** Snapshots of a local projection of a *pnr-gal4, UAS-EGFRdsRNA* pupal notum expressing E-cad-tdTomato and GC3Ai. The white stars show three neighbouring cells dying. Green, light and dark purple cell contours correspond to the curves shown in **e**. Scale bar=10μm. **e:** GC3Ai intensity in the three cells marked in **d**. Time 0 is the termination of extrusion of the first dying cell (green contour). Note that there is no inflexion of GC3Ai signal in the neighbours. **f:** Averaged GC3Ai differential signal (caspase activity) in the dying cell (green) and its neighbours (purple) upon depletion of EGFR in the notum. Light areas are s.e.m.. Time 0 is the termination of extrusion of the first dying cell. On average, caspase activity is maintained in the neighbouring cells (differential >0, compare with **Figure 3b,c**). **g:** Local projection of a E-cad-GFP pupae depleted for EGFR (*pnr-gal4, UAS-EGFR dsRNA*). Red dots show all the dying cells over the course of the movie (21 hours). Scale bar=10μm. The white rectangle show the analysis region (similar to the region used for the WT conditions). **h:** Averaged differences between the experimental distribution of cell death density in four EGFR depleted pupae and the corresponding simulated distribution (assuming independent events and the same death intensity). The inset shows the same analysis for WT pupae (from **Figure 2d**). Note the absence of yellow in the bottom left corner compared to the WT (no reduction of cell death density in EGFR depleted pupae for 10-60 min, <7 μm). **i:** Closest neighbour analysis of cell death distribution in EGFR depleted pupae. Integrated death probability for different time windows after cell elimination (20min-1h20, 1h20-2h20, 2h20-3h20) at different distances from the dying cells. While the first hour distribution was different in the WT (see top left inset, blue curve, from **Figure 2 e-g**), there is no more differences between curves upon EGFR depletion. **j:** Averaged map of dispersion p-value from the K-functions for 4 *pnr-gal4 EGFR RNAi* movies (pseudoclour: p-value, blue=no significant dispersion). The top right inset shows the averaged map of dispersion p-value for control movies (see **Figure 2i**) **k:** Number of occurrences of clustered elimination (>=3 cells) per movie (16 to 20 hours) in the WT pupae (5 nota) and upon depletion of EGFR (*pnr-gal4, UAS-EGFR dsRNA*, 9 pupae). **l:** Snapshots of E-cad-GFP local projection in a EGFR depleted pupae showing concomitant elimination of three cells (orange area) and an aberrant extrusion (relaxation at −30 and wound healing figure with E-cad accumulation at vertices, bottom left inset, compare with **Figure 1 c,d**). Time 0 is the termination of cell elimination. **m:** Evolution of the clone area shown in **l**. The grey zone corresponds to the relaxation phase and is followed by wound healing. Inset shows increased of clone solidity (area/convex area) during the relaxation and wound healing phases. **n:** Schematic of the local ERK feedback (red) and its impact on the distribution of cell elimination (green cells: caspase activation). Upon EGFR depletion, simultaneous caspase activation and cell elimination can occur, which leads to aberrant extrusion and transient loss of sealing (phase outlined in yellow, white area between green aggregates) followed by wound healing.

High rates of cell elimination are widespread during development and in adult tissues with high cell turnover. Similar local feedback mechanisms may be required for coherent cell elimination in the gut to prevent transient and recurrent sealing defects that could lead to chronic inflammation and inflammatory bowel disease^18^. Importantly, the mechanism we characterised in this study are conserved in mammals as similar ERK dynamics were observed near dying MCF10A cells which also generate transient resistance to apoptosis (personal communication, Olivier Pertz lab). Interestingly, other reports also described Ca^2+^ and ERK activation waves emanating from extruding MDCK and MCF10A pretumoral cells or caspase-induced extrusion^19,20^. However in these cases the waves rather promote extrusion of the dying/extruding cells through collective convergent movements. In contrast, we still see a high rate of cell extrusion upon depletion of ERK feedbacks in our tissue (*UAS-EGFR dsRNA*). Strikingly, the range of the communication seems to be very different between the notum (one cell row) and mammalian cells (10 to 15 cells away), suggesting either that the relays of communication are different, or that mechanical properties of these cells are very different.

Although at this stage we cannot rule out the contribution of contact-dependent communication for ERK activation in dying cell neighbours, several evidences point for a contribution of cell stretching. First, ERK activation dynamics correlate very well with transient cell stretching (**Fig. 3e,f**) and occurs very fast (few minutes). Moreover, we previously showed that cell stretching could promote cell survival through EGFR^10^. Accordingly, we found that EGFR depletion completely abolished ERK feedback near dying cells. We exclude the contribution of calcium signaling or an active secretion of the ligand from the dying cell, while laser induced wound healing was sufficient to promote neighbouring cell stretching and ERK activation. Other recent works have characterized the rapid effect of cell stretching on ERK activation^21,22^, which was recently shown to also rely on EGFR^23^. Further work characterising the molecular mechanism of cell deformation sensing by ERK will be required to fully test the contribution of mechanics in extrusion-induced ERK activation.

The occurrence of aberrant extrusion remarkably increases for clusters of three cells or more. However, we do not know at this stage what triggers these abnormal extrusions. On the one hand, abortive extrusion may be related to the distance that needs to be closed by the neighbouring cells: while this can be short for lines of cells (using the shortest axis), it will be higher for clusters and may prevent fast fusion of neighbouring cell junctions and gap closure^24^. On the other hand, abortive extrusion may be related to caspase-induced remodeling of tricellular junctions^25^, which are shared by caspase positive cells only in the case of clustered eliminations. Interestingly, some tricellular junction components have been shown to be cleaved by caspases^26^. Further characterisation of extrusion dynamics will be required to test these hypotheses.

Interestingly, 2D simulations of cell disappearance combined with transient death refractory phase in the neighbours suggest that ERK feedback could have a significant impact on the total number of dying cells (see **Figure S5a**, ~1.4 folds reduction for a 60 minutes refractory phase and for the rate of cell elimination we observed in the posterior region of the notum). This buffering effect increases sharply with the rate of cell elimination (**Figure S5a**). As such, it may be essential in conditions of high stress to dampen the rate of cell elimination and prevent tissue collapse. Moreover, this feedback could be sufficient to increase the rate of cell death near clones resistant to apoptosis when they are located in regions with a high rate of cell death (see **Figure S5b**, ~1.5 folds increase for cells completely surrounded by cells resistant to apoptosis in our conditions). This suggests that extrusion-driven ERK feedbacks may be sufficient to recapitulate some of the features of cell competition: namely a contact and context dependent increase of cell death near mutant clones^27^. Altogether, we propose that epithelial robustness and plasticity may be emerging features of local and transient ERK feedbacks driven by cell death.

## Acknowledgements

We thank members of RL lab for critical reading of the manuscript, especially Alexis Matamoro-Vidal for suggestions on the manuscript organisation. We would like to thank Jakub Voznica for initiating observations of miniCic in the pupal abdomen during his internship. We also thank Virgile Andreani for help in improving simulations, statistical analysis of the data, and some mathematical expressions. We are also grateful to Magalie Suzanne, the Bloomington Drosophila Stock Center, the Drosophila Genetic Resource Center, the Vienna Drosophila Resource Center for sharing stocks and reagents. We also thank B Aigouy for the Packing Analyser software and J. Ellenberg group for MyPic autofocus macro. LV is supported by a Post-doctoral grant “Aide au Retour en France” from the FRM (Fondation pour la Recherche Médicale, ARF20170938651) and a Marie Sklodowska-Curie postdoctoral fellowship (MechDeath, 789573), work in RL lab is supported by the Institut Pasteur (G5 starting package), the ERC starting grant CoSpaDD (Competition for Space in Development and Disease, grant number 758457), the Cercle FSER and the CNRS (UMR 3738).

## Authors contribution

RL and LV discussed and designed the project. RL did the experiments with GC3Ai and wrote the manuscript with LV. AD performed the simulations of the theoretical distribution of cell death, analysed experimental and simulated data using density maps and p-values maps, wrote theoretical expressions of death rates. LV and AD designed 2D numerical models to reproduce the effect of the feedback. FL designed miniCic-Scarlet flies and designed and performed the preliminary test of the optoDronc fly line. LV and MC set up Dextran injection in the pupal notum and initiated experiments with optoDronc. FCC provided the Spitz RNAi movies and part of the WT pupae movies. LV performed all the other analysis and experiments. Every author has commented and edited the manuscript.

## Methods

### Drosophila melanogaster husbandry

All the experiments were performed with *Drosophila melanogaster* fly lines with regular husbandry techniques. The fly food used contains agar agar (7.6 g/l), saccharose (53 g/l) dry yeast (48 g/l), maize flour (38.4 g/l), propionic acid (3.8 ml/l), Nipagin 10% (23.9 ml/l) all mixed in one liter of distilled water. Flies were raised at 25°C in plastic vials with a 12h/12h dark light cycle at 60% of moisture unless specified in the legends and in the table below (alternatively raised at 18°C or 29°C). Females and males were used without distinction for all the experiments. We did not determine the health/immune status of pupae, adults, embryos and larvae, they were not involved in previous procedures, and they were all drug and test naïve.

### Drosophila melanogaster strains

The strains used in this study and their origin are listed in the table below.

| Fly line | Chromosome location | Origin (citation) | Source |
| --- | --- | --- | --- |
| *E-cad-tdTomato (KI)* | II | <sup>28</sup> | BDSC_58789 |
| *E-cad-GFP (KI)* | II | <sup>28</sup> | BDSC_60584 |
| *UAS-optoDronc* | II (attp40) | This study | This study |
| *Tub-miniCic-mScarlet* | II | This study | This study |
| *ubi-EKARNls* | II | <sup>16</sup> | Shigeo Hayashi |
| *UAS-EGFR dsRNA* | II | VDRC | KK 107130 |
| *UAS-spitz dsRNA* | II | VDRC | GD 3922 |
| *UAS-GcAMPx20* | III | Bloomington | BDSC_32236 |
| *UAS-GC3Ai* | III | <sup>12</sup> | Magalie Suzanne |
| *GMR-gal4* | II | Bloomington | BDSC_1104 |
| *UAS-his3.3 mIFP-T2A-H01* | III | Bloomington | BDSC_64184 |
| *Hs-flp22; ; Act<cd2<gal4,UAS-GFP* | I; III | <sup>10</sup> | <sup>10</sup> |

The exact genotype used for each experiment is listed in the next table. ACI: time After Clone Induction, APF: After Pupal Formation, n: number of pupae/adults.

| Figure | Genotype | Heat shock 37°C | Stage | n |
| --- | --- | --- | --- | --- |
| 1b-j | *hs-flp22; E-cad-tdTomato (KI)/UAS-optoDronc ; act<y+<gal4 , UAS-GFP/+* | 12 min, 2 days ACI | 30h APF | 10 |
| 2a | *E-cad-GFP (KI), tub-miniCic-mScarlet ; pnr-gal4, UAS-His3-mIFP* | N/A | 16h APF | 5 |
| 3a-c | *E-cad-tdTomato (KI)/+; pnr-gal4/UAS-GC3Ai* | N/A | 20h APF | 3 |
| 3d-f | *E-cad-GFP (KI), tub-miniCic-mScarlet ; pnr-gal4, UAS-His3-mIFP* | N/A | 20h APF | 5 (quantified on 2) |
| 3g | *E-cad-GFP (KI), tub-miniCic-mScarlet/UAS-EGFR dsRNA ; pnr-gal4, UAS-His3-mIFP/+* | N/A | 16h APF | 3 (quantified on 2) |
| 4a-c | *tub-miniCic-mScarlet ; pnr-gal4/UAS-GC3Ai* | N/A | 20h APF | 3 |
| 4d-f | *E-cad-tdTomato (KI)/UAS-EGFR dsRNA; pnr-gal4/UAS-GC3Ai* | N/A | 20h APF | 3 |
| 4g-l | *E-cad-GFP (KI), tub-miniCic-mScarlet/UAS-EGFR dsRNA ; pnr-gal4, UAS-His3-mIFP/+* | N/A | 16h APF | 9 |
| S1a | *E-cad-tdTomato (KI)/UAS-optoDronc; pnr-gal4/+* | N/A | 16h APF | 2 |
| S1b | *GMR-gal4/UAS-optoDronc (dark)* | N/A | Adult | 20 |
| S1b | *GMR-gal4/UAS-optoDronc (blue light)* | N/A | Adult | 19 |
| S1b | *GMR-gal4/UAS-optoDronc ; UAS-p35/+ (blue light)* | N/A | Adult | 46 |
| S1c | *hs-flp22; E-cad-tdTomato (KI)/UAS-optoDronc ; act<y+<gal4 , UAS-GFP/+* | 12 min, 2 days ACI | 30h APF | 3 |
| S1c | *hs-flp22, E-cad-tdTomato (KI)/UAS-optoDronc ; act<y+<gal4 , UAS-GFP/UAS-p35* | 12 min, 2 days ACI | 30h APF | 3 |
| S1f | *hs-flp22, E-cad-tdTomato (KI)/UAS-optoDronc ; act<y+<gal4 , UAS-GFP/+* | 12 min, 2 days ACI | 30h APF | 3 |
| S1f | *E-cad-GFP (KI), tub-miniCic-mScarlet ; pnr-gal4, UAS-His3-mIFP* | N/A | 30h APF | 5 |
| S2a | *E-cad-GFP (KI), tub-miniCic-mScarlet ; pnr-gal4, UAS-His3-mIFP* | N/A | 16h APF | 5 |
| S2b | *E-cad-GFP (KI), tub-miniCic-mScarlet/ UAS-EGFR dsRNA ; pnr-gal4, UAS-His3-mIFP* | N/A | 16h APF | 4 |
| S3a,b | *tub-miniCic-mScarlet ; pnr-gal4/ubi-EKARNls* | N/A | 20h APF | 4 |
| S3c,d | *E-cad-GFP (KI), tub-miniCic-mScarlet* | N/A | abdomen (20h APF) | 2 |
| S3e | *Hs-flp22; E-cad-GFP (KI), tub-miniCic-mScarlet/UAS-EGFR dsRNA ; act<y+<gal4, UAS-His3-mIFP/+* | 20 min, 2 days ACI | 20h APF | 3 |
| S4a,b | *E-cad-tdTomato (KI)/+; pnr-gal4/UAS-GCaMPx20* | N/A | 16h APF | 2 |
| S4c,d | *Hs-flp22; E-cad-GFP (KI), tub-miniCic-mScarlet/UAS-spitz dsRNA ; act<y+<gal4, UAS-His3-mIFP/+* | 20 min, 2 days ACI | 20h APF | 2 |
| S4e,f | *E-cad-GFP (KI), tub-miniCic-mScarlet ; pnr-gal4, UAS-His3-mIFP* | N/A | 20h APF | 7 |

### Design of optoDronc

The GFP-linker-DRONC-linker-CRY2PHR was first cloned in the pCaspeR4-Tubp-Gal80 vector (addgene 17748) and subsequently cloned in the pJFRC4-3XUAS-IVS-mCD8::GFP (Addgene 26217). Initial cloning was performed by three successive amplifications/ligations. Briefly, GFP was first inserted in pCaspeR4-Tubp by PCR-amplifying GFP from pJFRC19-13XLexAop2-IVS-myr::GFP (Addgene 26224) adding NotI, BglII, XbaI restriction sites and linkers, and eventually ligation into pCasper4-TubP-Gal80 after digestion with NotI and XbaI. Dronc cDNA was then inserted in this plasmid through PCR amplification on the cDNA clone LP09975 (DGRC) adding BglII, NheI and XbaI restriction sites and linkers, and then ligation into pCasper4-TubP-GFP-linker cut with BglII and XbaI. Finally, CRY2PHR was inserted by amplifying residues 1-498 of CRY2 from pGal4BD-CRY2 (Addgene 28243) while adding NheI and XbaI sites and ligation into pCasper4-TubP-GFP-linker-DRONC-linker cut with NheI and XbaI. The GFP-linker-Dronc-linker-CRY2PHR was cut with NotI and XbaI and inserted by ligation in pJFRC4-3XUAS-IVS-mCD8::GFP (Addgene 26217) after digestion with NotI and XbaI. The construct was checked by sequencing and inserted at the attp site attp40A after injection by Bestgene. The primers used for the construct are listed below (restriction sites in bold and linker in italic).

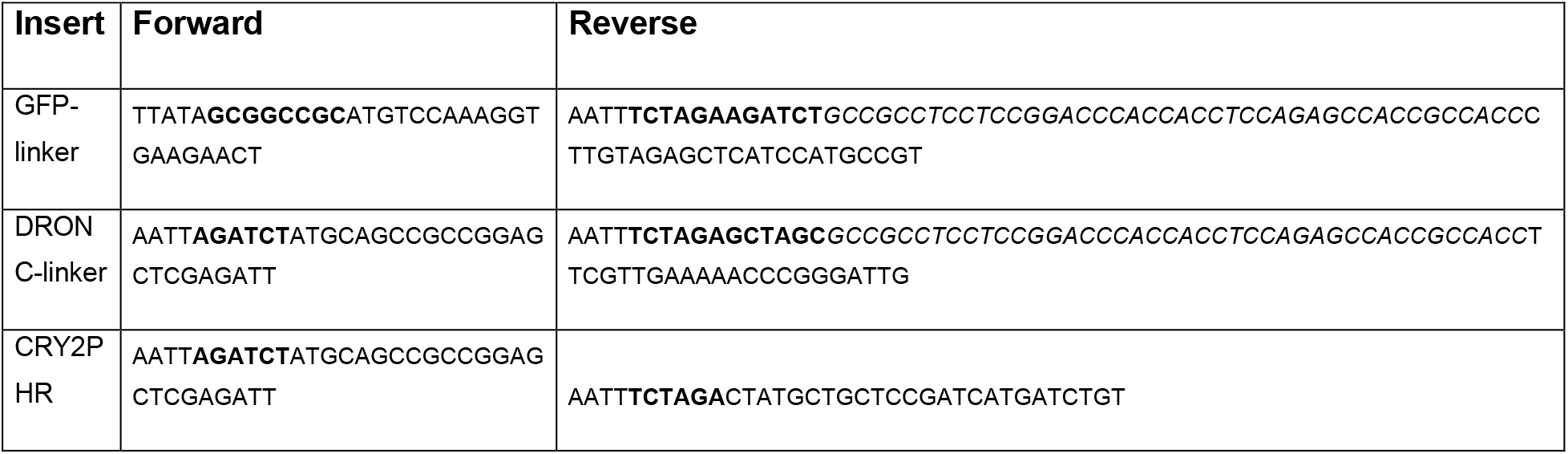

### Induction of cell death using optoDronc

To induce optoDronc in the eye, GMR-gal4 females were crossed with homozygous males UAS-optoDronc. Tubes containing crosses and progeny were either kept in the dark at 25°C (control) or maintained in a cardboard box permanently lighted by a blue LED array (LIU470A, Thorlab) in the same 25° incubator as the control. Female adult eyes were then imaged on a Zeiss stereoV8 binocular equipped with a colour camera (Axiocam Icc5).

For induction of optoDronc in clones in the pupal notum, *hs-flp; E-cad-tdTomato(KI); act<cd2<G4* females were crossed with homozygous *UAS-optoDronc* or *UAS-optoDronc; UAS-p35*. Clones were induced through a 12 minutes heat shock in a 37°C waterbath. Tubes were then maintained in the dark at 25°C. White pupae were collected 48 hours after clone induction and aged for 24h at 25°C in the dark. Collection of pupae and dissection were performed on a binocular with LED covered with home-made red filter (Lee colour filter set, primary red) after checking that blue light was effectively cut (using a spectrometer). Pupae were then imaged on a spinning disc confocal (Gataca system). The full tissue was exposed to blue light using the diode 488 of the spinning disc system (12% AOTF, 200ms exposure per plane, 1 stack/min). The proportion of remaining cells was calculated by measuring the proportion of cells remaining in the tissue at time t compared to t0. Extrusion profiles were obtained by segmenting extruding cells in the optoDronc clones with E-cad-tdTomato signal (only single cell clones in this case) or WT cells marked with E-cad-GFP in the posterior region of the notum using Packing analyser^29^. Curves were aligned on the termination of extrusion (no more apical area visible) and normalised with the averaged area on the first five points. Clone area profile (**Figure 1d, Figure 4m**) were obtained by segmenting the group of cells in the clone and calculating the area and the solidity (Area/Convex Area) on Matlab. Categorisation of normal versus abnormal extrusion was based on the dynamics of clone contraction: transient relaxation or accumulation of E-cad at vertices was counted as abnormal extrusion. Large relaxation combined with clear E-cad accumulation at vertices was counted as “transient holes”.

### Design of miniCic-mScarlet

The pCaspeR4-Tubp-miniCic-linker-mScarlet-I was obtained by amplifying by PCR miniCic-linker from pCaspeR4-Tubp-miniCic-linker-mCherry^10^ and mScarlet-I from pmScarlet-i_C1 (Addgene 85044). These two inserts were cloned in the vector pCaspeR4-TubP-Gal80 linearised by NotI, XbaI digestion (to excise Gal80) using NEBuilder HiFi DNA Assembly Method. The construct was checked by sequencing and inserted through P-element after injection by Bestgene. The primers used for the construct are listed below.

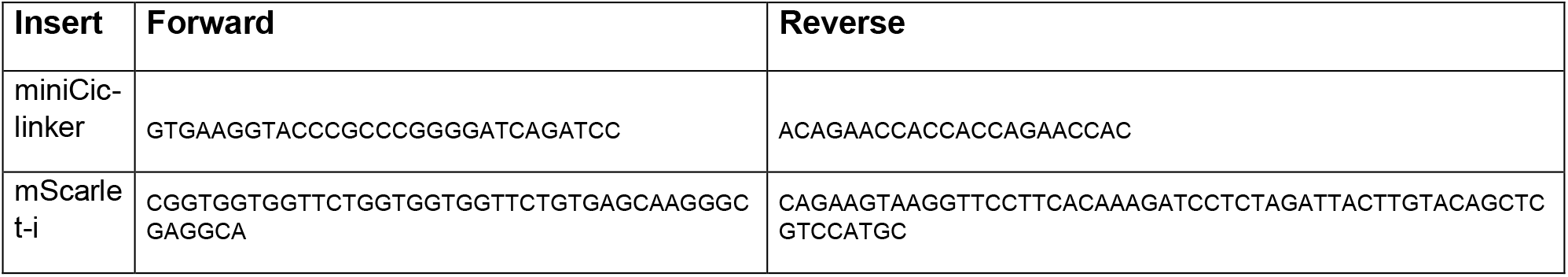

The construct was tested by comparing the dynamics and pattern in the notum compared to previously characterised dynamics using miniCic-mCherry^10^. Similar dynamics were also observed at the single cell level between miniCic-mScarlet and the FRET sensor ubi-EKARnls^16^ (**Fig. S3**).

### Live imaging and movie preparation

Notum live imaging was performed as followed: the pupae were collected at the white stage (0 hour after pupal formation), aged at 29°, glued on double sided tape on a slide and surrounded by two home-made steel spacers (thickness: 0.64 mm, width 20×20mm). The pupal case was opened up to the abdomen using forceps and mounted with a 20×40mm #1.5 coverslip where we buttered halocarbon oil 10S. The coverslip was then attached to spacers and the slide with two pieces of tape. Pupae were collected 48 or 72h after clone induction and dissected usually at 16 to 18 hours APF (after pupal formation). The time of imaging for each experiment is provided in the table above. Pupae were dissected and imaged on a confocal spinning disc microscope (Gataca systems) with a 40X oil objective (Nikon plan fluor, N.A. 1.30) or 100X oil objective (Nikon plan fluor A N.A. 1.30) or a LSM880 equipped with a fast Airyscan using an oil 40X objective (N.A. 1.3), Z-stacks (1 μm/slice), every 5min using autofocus at 25°C. The autofocus was performed using E-cad-GFP plane as a reference (using a Zen Macro developed by Jan Ellenberg laboratory, MyPic) or a custom made Metamorph journal on the spinning disc. Movies were performed in the nota close to the scutellum region containing the midline and the aDC and pDC macrochaetae. Movies shown are adaptive local Z-projections. Briefly, E-cad plane was used as a reference to locate the plane of interest on sub windows (using the maximum in Z of average intensity or the maximum of the standard deviation). Nuclear signal was then obtained by projecting maximum of intensity on 7 μm (7 slides) around a focal point which was located 6 μm basal to adherens junctions (see ^10^ for more details).

### Laser ablation

Photo-ablation experiments were performed using a pulse UV-laser (355nm, Teem photonics, 20kHz, peak power 0.7kW) coupled to a Ilas-pulse module (Gataca-systems) attached to our spinning disk microscope. The module was first calibrated and then set to 40-60% laser power. Images were taken every 500ms and 3 to 6 single cells were ablated 10 images after the beginning of the movie (20 repetitions of “point” ablation ~50ms exposure per cell). Cells were selected in regions with high and homogeneous nuclear miniCic levels. For each single cell ablation, miniCic signals over time were extracted in 4 cell nuclei of the first row and second row of cells around the ablated cell. Nuclei were visualized and clicked by hand using the Histone3-mIFP fluorescent channel after a local z-projection. Background value for each movies (for miniCic) were extracted and removed from the signals, then the single cell miniCic signal was normalized to 1 using the first time point.

### Dextran injection in pupal notum

30h APF pupae were glued on double sided tape and the notum was dissected with red filter. Pupae were then injected using home-made needles (borosilicate glass capillary, outer diameter 1mm, internal diameter 0.5mm, 10cm long, Sutter instruments) pulled with a Sutter instrument P1000 pipette pulling apparatus. Dextran Alexa 647 10,000 MW (Thermofisher, D22914) was injected at 2mg/ml in the thorax of the pupae using a Narishige IM400 injector using a constant pressure differential (continuous leakage) and depressurisation in between pupae. Imaging of Dextran leakage in live pupae was performed after local projection using E-cad plane as a reference to measure Dextran concentration at the junction plane (note that septate junctions are located basally to adherens junctions). Dextran intensity was measured using a ROI in the center of the clone after normalization using the first 5 time points and removal of intensity background (estimated on a region with no signal. Note that Dextran injection cannot be used to track leakages over long time scales (>4 hours) as the majority of Dextran get rapidly trapped in fat body cells and in epithelial cell endosomes.

### Signaling dynamics and cross-correlation

#### Measurement of ERK activity

ERK dynamics were measured using the mean nuclear intensity of miniCic. Whenever possible, a marker of the nucleus (His3-mIFP) was used to track the nucleus manually (Fiji macro). EKAR FRET signal was obtained after local projection on 6 planes of CFP and YFP signals. CFP and YFP signals were blurred (Gaussian blur, 2 pixel width) and the ratio of YFP/CFP signal was then calculated. Raw YFP-nls signal was thresholded and used as a mask to only keep nuclear FRET signal. For the comparison of miniCic and EKAR, single curves from the same cell were aligned on the peak of the FRET signal (maximum value) and eventually averaged for all the cells.

For the measurement of miniCic in neighbours combined with apical area (**Figure 3**), cell contours were tracked using E-cad-GFP (Packing analyser), while nuclei were manually tracked on Matlab using His3-mIFP. For each dying cell, the nuclear miniCic intensity and apical area were averaged for all the neighbours (first row and second row of cells). The final curves are averaged of each of these curves after alignment at the termination of cell extrusion (apical area=0). Note that the “n” used for s.e.m. calculation is not the total number of neighbours, but the number of cell clusters.

#### Analysis of GC3Ai signal

To analyse caspase activity, we used the differential of GC3Ai signal as a proxy. GC3Ai becomes fluorescent upon cleavage of a domain by effector caspases which triggers GFP folding and maturation^12^. The dynamics of GFP signal can be written as follows:

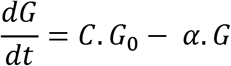

Where G stands for concentration of active GFP, C concentration of active effector caspase, G_0_ concentration of inactive GFP, and α rate of GFP degradation.

On timescales of one hour, we can neglect GFP degradation. Assuming a constant pool of inactive GC3Ai, G_0_ can be considered as a constant. As such:

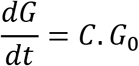

Therefore the change of intensity over time should be proportional to effector caspase activity (C). Practically, GC3Ai signal was obtained after local projection (using E-cad or nuclear miniCic as a reference) and measurement of mean intensity on small circular ROI at the center of the cell (Fiji). The differential was calculated using the Diff function of Matlab after smoothing the intensity curve (10 time points averaging window) and aligning the curve on the time of disappearance (end of extrusion) of the first cell. Note that the analysis of caspase reversion was restricted to neighbours having a positive slope before the elimination of the first cell (in order to see caspase shutdown).

To cross-correlate GC3Ai and ERK dynamics, local projections centered on nuclear miniCic were used (with GC3Ai plane of interest shifted apically). miniCic nuclear intensity and GC3Ai cytoplasmic intensity were measured by manually tracking small circular ROI (20px) in the center of the cells (Fiji home-made macro). The cross-correlation between GC3Ai differential and miniCic nuclear signal was calculated on Matlab with the “xcorr” function with the ‘coef’ option (normalised cross-correlation). All the curves (one per cell) were then averaged.

#### Measurement of GCaMP

GCaMP signal was measured on local projections using E-cad-tdTomato as a reference plane after global correction for bleaching. Each cell extrusion was cropped and the movie realigned using the Stackreg Fiji plugin. GCaMP signal was then measured using the mean intensity in a 20×20 pixel square in the center of each cells. Each single cell curve was aligned (time 0 at the termination of extrusion) and normalised using 5 time values around time 0.

### Statistical analysis of cell death distribution

#### Calculation of the probability to observe three cells clustered elimination

We wanted to evaluate the probability to observe a cluster of three cells eliminated in less than 30 minutes. We first considered a hexagonal grid of points in a squared array (N cells per side) and assumed periodic boundaries. The probability of cell disappearance over 30 min is p. As such the probability to observe a specific cluster of 3 cells eliminated in the same 30 min window is p^3^. The total number of possible 3 cells configurations is approximatively given by: 2.*N*^2^.

Using the posterior region of WT nota, we estimate p~0.04 (probability of cell elimination in 30 minutes). As such for a group of 400 cells (region of interest used in **Figure 2**) observed for 20 hours (T= 40 times 30 minutes), the expected number of 3 cell-cluster elimination is given by: *E* (*number of clusters*) = 2*N*^2^*p*^3^*T* = 2.05.

Therefore, we would expect to observe several occurrences of 3 cell-cluster eliminations per movie.

#### Analysis of death distribution

For the analysis of death distribution we selected movies lasting 16 to 20 hours in the posterior region of the pupal notum. Cell extrusion events were clicked by hand and checked twice. Extrusion localisations were corrected for global drift (translation) of the tissue (Matlab procedure). Spatio-temporal distribution of death was first checked by x-y-t histogram. According to that distribution, for each movie, an area of interest (in time and space) was selected by hand to obtain an area where distribution was as homogeneous as possible (double checked by x-y-t histograms). These data sets were then the basis of three statistical analyses (see below).

##### 1. Distribution of distances

Distances in time and space between all couples of extrusions were computed. For each data set we plotted the maps of densities of death at a given distance (number of deaths divided by the area of the disc considered) as a function of spatial and temporal distances from each dying cell. From each data set we estimated the effective death rate, i.e. the intensity of the random spatio-temporal process, and used it to simulate the corresponding Poisson process 200 times. For each simulation we performed the same analysis as for the experimental data set, namely calculating maps of death densities,and eventually averaged the 200 maps. Finally, for each movie we calculated the difference between the averaged simulated map and the corresponding experimental map. We show in the main figures the average of the “difference map” for every experiment (5 for WT, 4 for EGFR-RNAi).

##### 2. Closest neighbour analysis

For each death event, we calculated the Euclidean distance to the closest death in a given time window. Based on the previous analysis, we selected windows of 20’ to 1h20’, 1h20’ to 2h20’ and 2h20’ to 3h20’. We excluded the first 20 minutes as they correspond to the characteristic time of extrusion (where cells cannot be reverted anymore). We then ordered these shortest distances by size and plotted the cumulative probability of closest death at a given distance. A value of 40% at a distance of 10 micrometers means than 40% of the closest deaths are localized between 0 and 10 micrometers away.

##### 3. p-values test using “K-functions” and random labeling test

This procedure is used for estimating how far is one spatio-temporal process from a purely random Poisson process^11^. The test is performed in two main steps. The first step consists of calculating so-called “K-functions” of the given process, and the second is the random re-labeling test as explained below.

The space-time K-function of the observed process, *K*^0^(*r, h*), is defined as the expected number of additional events within the space-time distance (*r, h*) of a randomly selected event. It depends on the intensity *λ* of the spatio-temporal process and it can be estimated using only the observed events and without additional assumption on the process.

The complete absence of dispersion or clustering in a given process implies that there is no relation between the location and timing of events. This is formally given through the temporal indistinguishability hypothesis. Under this hypothesis, there is an equal probability to observe our set of events, or events with the same locations in space but with shuffled times. From here we can perform the random labeling test as follows.

We draw a sample of *N* random permutations of the times of events. For a given space-time distance (*r, h*) we calculate the value of the K-function, *K*(*r, h*), for each permutation. We compute the number *M*(*r, h*) of sampled re-labelings with the value of the K-function smaller than the one of the original labeling, i.e. *K*(*r, h*) < *K*^0^(*r, h*). Then, the probability of obtaining a value as small as *K*^0^(*r, h*) is estimated by the space-time dispersion p-value: *p*(*r, h*) = (*M*(*r, h*) + 1)/(*N* + 1). If we repeat the same procedure for all space-time distances (*r, h*), we obtain the map of dispersion p-values.

This procedure is implemented in Python. The maximum distance is set to *r_max_* = 25*μm*, and the maximum temporal interval to *h_max_* = 150*min*. We subdivided these maximum values in 25 and 30 equal increments (1*μm* and 5*min*). We simulated 9999 re-labelings of times to test for space-time dispersion of the extrusion events. The results of these 750 tests are plotted on a grid and then interpolated to obtain a p-value map. We performed this procedure for each movie.

### Calculation of the effective death rate with caspase resistant neighbours

We assume that the cell has six neighbours, and *n* neighbours resistant to apoptosis (*n* ≤ 6), and that the death of one neighbour will protect the cell for a period *T*. The probability of a cell to die at the time *t* is the probability that none of its (6 − *n*) sensitive neighbours died during the period *T* prior to *t*, multiplied by its intrinsic probability to die at the moment *t*. The death of each isolated sensitive cell is a random event modelled as a temporal Poisson process, with the event rate *λ* (expected number of cell deaths per minutes).

By definition, for the Poisson process *X* with rate *λ* the probability of the event (death) to happen in the time interval [*t, t + dt*], for *dt* very small, is:

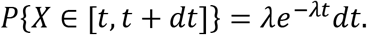

From here, the probability that an isolated sensitive cell dies during a time period of length *T* is:

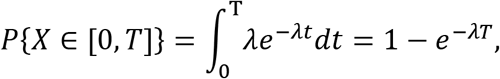

and the probability that it does not die during this time is: 1 − (1 − *e^−λT^*) = *e^−λT^*.

The probability that the cell dies in a given moment *t* is defined as the probability to die in the time interval [*t, t + dt*], for *dt* very small, under assumption that it didn’t die until the time *t*. It is the same as the probability that it dies in the time interval [0, *dt*]:

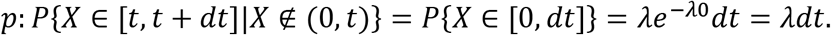

This is also one of the definitions for the rate of the temporal Poisson process.

Thus, for the cell with six neighbours, where *n* neighbours are resistant to apoptosis (*n* ≤ 6), the probability to die at the given moment *t* is:

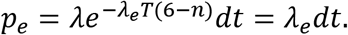

Here we approximated the self-inhibiting random process with a Poisson process with a different death rate, that we name the effective rate of death, and we used this rate to express the probability that the neighbouring cells do not die during the period *T*. Finally, the effective rate is:

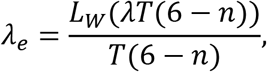

where *L_W_* is Lambert W function. Note that we neglect here the effect of the delay of inhibition on the neighbours. We used this expression in **Figure S5b**.

### Calculation of the intrinsic death intensity for the self-inhibiting spatio-temporal process, with continuous time and space

Assume that *X* is the spatio-temporal Poisson process. Then, by definition, the probability that the number of events that fall in the spatial region *C* during the time period [0, *t*] is equal to *k*, is:

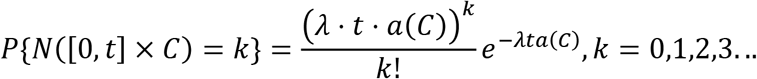

where *λ* is the space-time intensity of the process, i.e. it is the expected number of events per unit of space-time volume (expected number of deaths per minute per micrometer squared), and *a*(*C*) is the surface of the given region *C*.

Hence, the probability that no event (*k* = 0) happens on a given region *C* during a period of time *τ* is: *e*^−*λτa*(*C*)^. If the region *C* is a disk of radius *r*, its surface is *r*^2^*π*, and this probability is: *e*^−*λτr*^2^*π*^.

By definition of the space-time intensity of the process, the probability that the event happens in a given moment *t* and that it falls on a position (*x,y*), is given by:

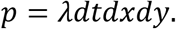

Now we consider the self-inhibiting spatio-temporal process. Meaning, for each event there is a feedback that inhibits the process on a disk around the event of radius *r*, and during a period of time *τ*. Such a self-inhibiting process is not anymore a Poisson process, however it can be approximated as one with a different effective intensity, *λ_e_*. The probability that the death happens at some time *t* and on a given location (*x,y*), is the probability that there was no event in its spatio-temporal vicinity (characterised by the spatio-temporal volume of the feedback), multiplied with the probability of an isolated cell death at this moment and place, or

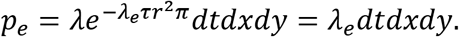

Then the observed or effective intensity, *λ_e_*, can be obtained by solving the equation:

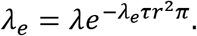

Finally,

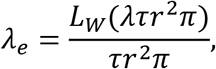

where *L_w_* is Lambert *W* function. We can also obtain an expression for the intrinsic intensity:

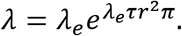

Note that with this approximation we neglected an effect of the delay in the feedback. This means that if we can estimate parameters of the feedback *τ* and *r*, and using the observed intensity, *λ_e_*, we can have an estimate on the intrinsic intensity of the spatio-temporal self-inhibiting process.

To verify this expression numerically, we perform the following test: assume we want to observe an effective intensity of the process *λ_e_* = 5.14 · 10^−5^*min*^−1^*m*^−2^ with the self-inhibiting process whose feedback has the parameters: *τ* = 40*min, r* = 5*μm*, and delay 10*min*. We first calculate, using the above expression the intrinsic rate of the process to be *λ* = 6.04 · 10^−5^. Using this rate and given parameters of the feedback, we run a long simulation (2 · 10^6^*min*) of the self-inhibiting process. We observed the effective intensity of this process to be: *λ_e_* = 5.12 · 10^−5^*min*^−1^*m*^−2^. For the comparison, we run the same length of the simulation of the actual Poisson process (without feedback) with the intensity *λ = λ_e_* = 5.14 · 10^−5^*min*^−1^*m*^−2^. The observed intensity of this simulated process is *λ* = 5.13 · 10^−5^*min*^−1^*m*^−2^. This confirms that we have a good approximation for expression of the intrinsic intensity of the self-inhibiting process. We performed such comparisons for several values of intrinsic intensity.

We use this expression to estimate an intrinsic intensity of the processes of the experimental data, and for the simulations of self-inhibiting processes that correspond to our experiments (**Figure 2h**)

### 2D simulations of death rate

To evaluate the effect of the duration of the feedback as a function of the intrinsic rate of death we made Python routine as follows:

We use a hexagonal grid as an approximation of the cell arrangement in the tissue. Each node in the grid represents a cell, with the intrinsic rate of death *λ*. Number of cells per side is *n*, so in total we have *n*^2^ cells. The cells can be in 2 different states: “regular” (sensitive to death), or “inhibited for death” (probability to die is 0). When one cell dies it inhibits itself, and its 6 neighbours to die during some time *T* (in minutes).

Each death, *e* = (*t_e_,x_e_,y_e_*), is described with the time of death, *t_e_*, and its coordinates in the grid (*x_e_,y_e_*). We simulate the times of death for the Poisson process with rate *l = λ · n*^2^. For each simulated time, *t_e_*, we assign a random position (*x_e_,y_e_*) in the grid. Then we check if any of its neighbours, or itself, has died in the previous *T* minutes. If there was no death recorded in its neighbourhood, the cell is a regular cell and we record a new death event (*t_e_,x_e_,y_e_*). If there was a death in its neighbourhood in the last *T* minutes the cell is refractory to death and we do not record this event.

At the end of simulation we compute the effective death rate as 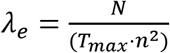, where *N* is a total number of recorded deaths and *T_max_* is the simulation time.

The parameters that we used in our simulations are the following: *n* = 25 (625 cells), *T_max_* = 5000*min*. We simulated effects of different inhibition times *T* ∈ [0,5,10,20,…,150]*min* with different intrinsic death rates, and repeated it 5 times per value. (**Figure S5a**)

### Statistics

Data were not analysed blindly. No specific method was used to predetermine the number of samples. The definition of n and the number of samples is given in each figure legend and in the table of the Experimental model section. Error bars are standard error of the mean (s.e.m.). p-values are calculated through t-test if the data passed normality test (Shapiro-Wilk test), or Mann-Whitney test/Rank sum test if the distribution was not normal. Statistical tests were performed on Graphpad Prism 8 or Matlab.

---

**Figure S1:**
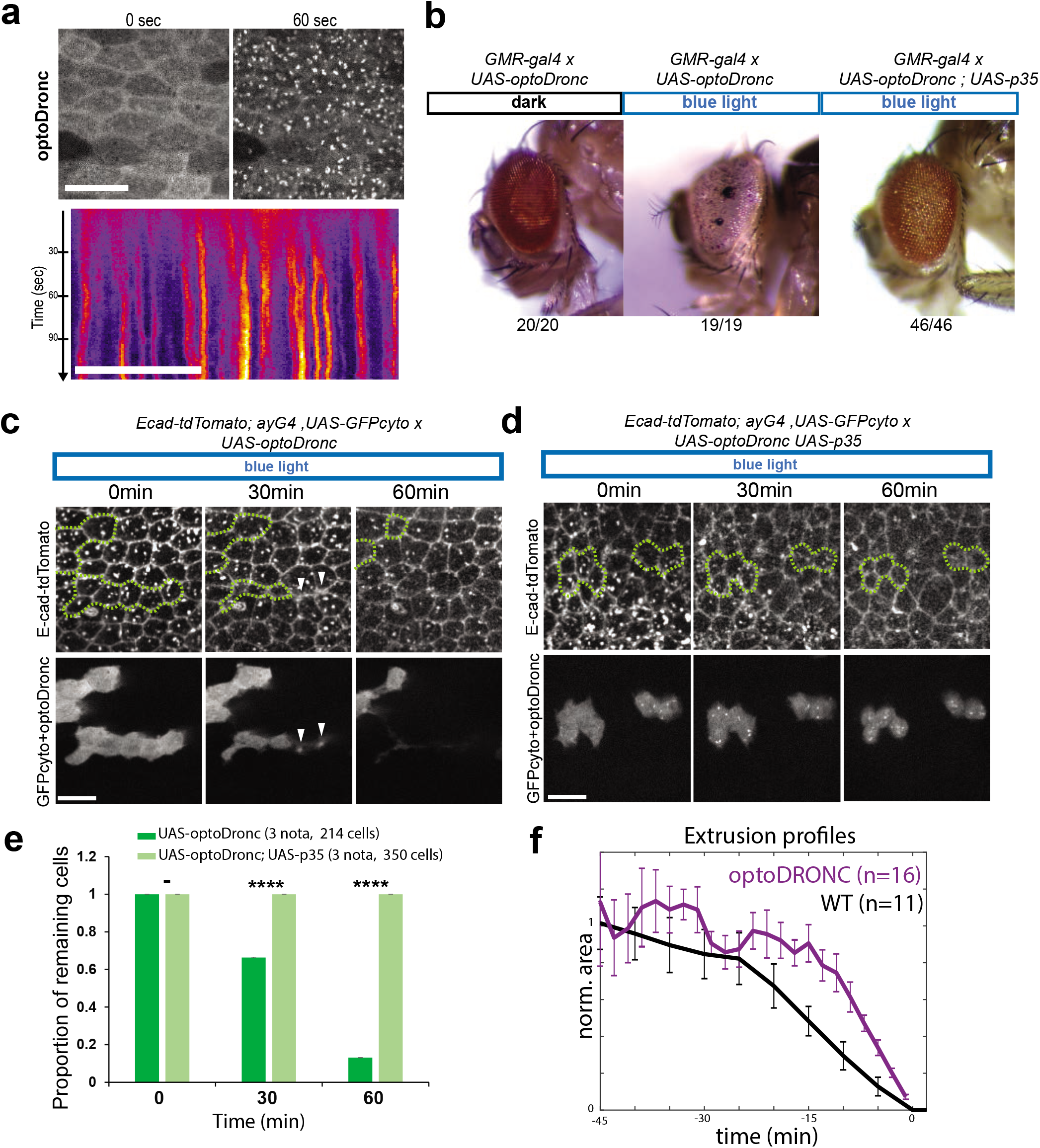
optoDronc triggers cell extrusion through effector caspase activation. **a:** Rapid clustering of UAS-optoDronc (GFP fusion) upon blue light exposure (single plane movie, 1 frame/sec). Scale bar= 10μm. Bottom: kymograph of optoDronc-GFP signal (clusters already appear after 20 sec) **b:** GMR-gal4 adult *Drosophila* eyes expressing UAS-optoDronc from flies raised in a blue light chamber (left), in the dark (middle) or with p35 overexpression (effector caspase inhibitor) in the light chamber. N are numbers of females. **c,d:** Snapshots of pupal nota (local projection) expressing *UAS-optoDronc* (**c**) or *UAS-optoDronc* and *UAS-p35* (**d**) in clones (bottom, UAS-GFP and optoDronc-GFP in the same channel) and E-cad-tdTomato. Green lines show clone contours. White arrowheads show extruding cells. Scale bars=10μm. **e:** Proportion of clonal cells remaining in the epithelium upon blue light exposure expressing *UAS-optoDronc* (green) or *UAS-optoDronc; UAS-p35* (light green). Error bars are 95% confidence interval,-= non-significant, ****=p<10^−4^. **f:** Averaged and normalised cell apical area during spontaneous extrusion (black curve, WT movie, frame rate 5min) or upon induction of extrusion through light activation of optoDronc in single cell clones (purple curve). Curves are aligned at time 0 min (cell apical area = 0). Error bars are s.e.m.. Note that optoDronc-triggered extrusion is slightly faster than WT extrusion but globally similar.

**Figure S2:**
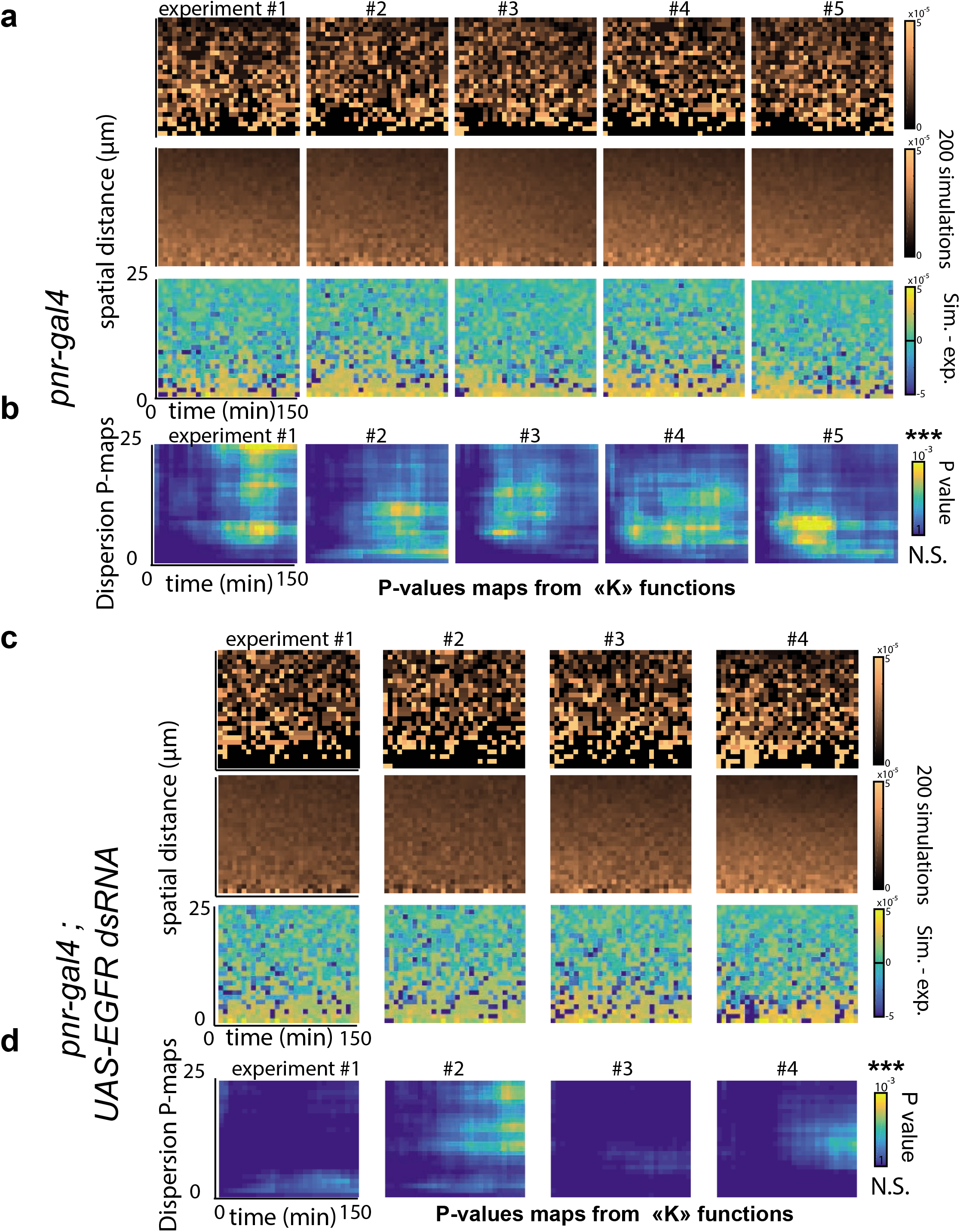
Distribution of cell death in control nota and upon depletion of EGFR. **a,c:** Maps of the local death density near each dying cells at different times (see **Figure 2b-d**) for control pupae (**a**, *pnr-gal4*) or upon depletion of EGFR (**c**, *pnr-gal4; UAS-EGFR dsRNA*). Top: experimental distribution (one map = one pupa), y-axis: spatial distance in μm, x-axis: time distance in minutes. Middle: simulations of the death distribution for the corresponding experiment assuming a Poisson process with the same effective death intensity (average of 200 simulations). Bottom: Difference between experiment and simulation distributions (yellow= low proportion of cell death compared to simulations). **b,d:** Maps of the dispersion p-value calculated using K-functions for control pupae (**b**, *pnr-gal4*) or upon EGFR depletion (**d**, *pnr-gal4; UAS-EGFR dsRNA*). y-axis: spatial distance in μm, x-axis: time distance in minutes. Pseudo-colours show the significance of the dispersion (yellow: significant dispersion, blue: no significant dispersion).

**Figure S3:**
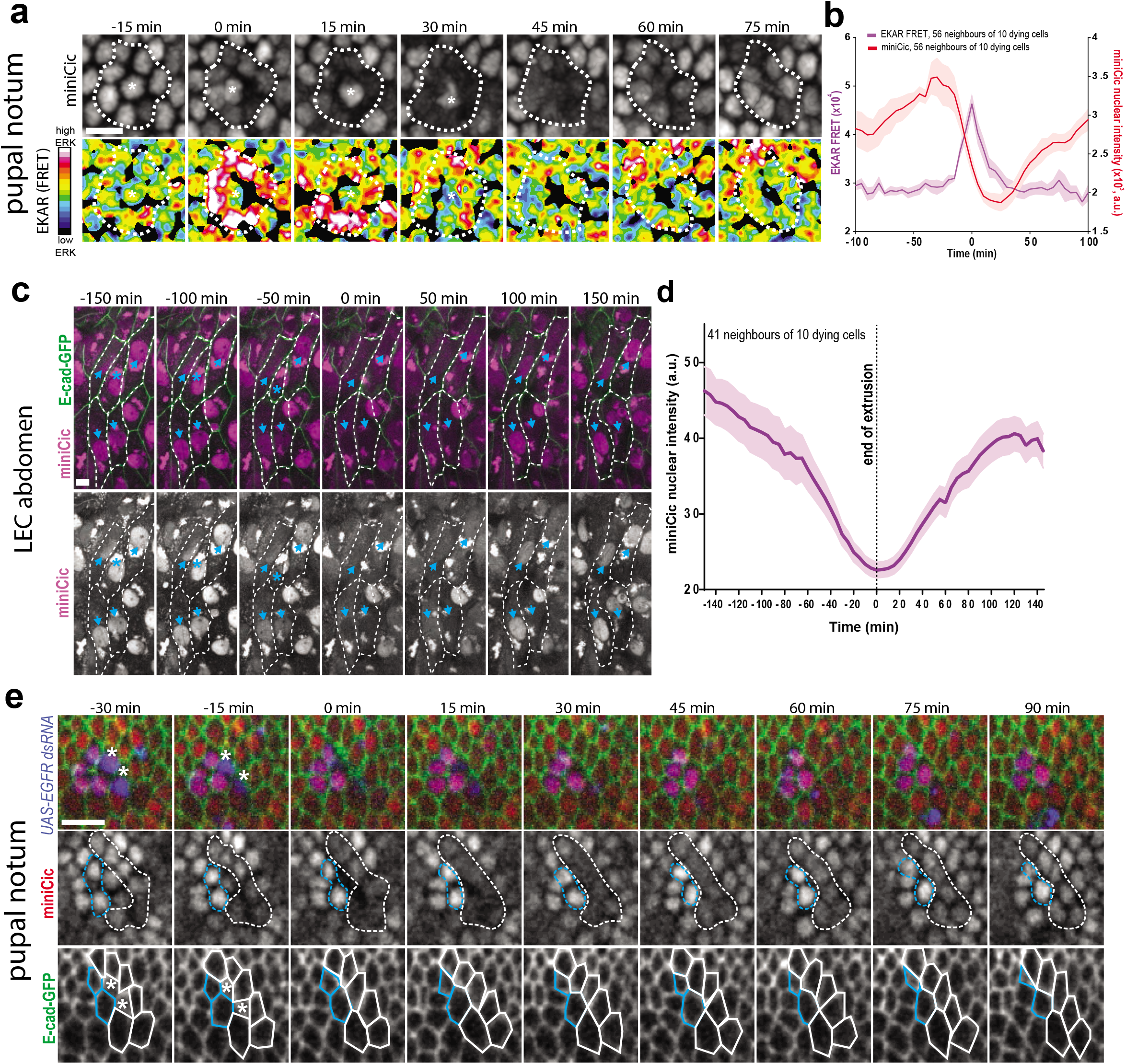
ERK pulses are also present in the abdomen and require EGFR cell-autonomously. **a:** Snapshots of local projections of a pupal notum expressing miniCic (top, grey) and a FRET sensor of ERK (EKAR, bottom). Red signal: high ERK, blue signal: low ERK. The white star marks a dying cell. The dotted lines show the miniCic and the FRET signal in the neighbouring cells. Scale bar=10μm. **b:** Averaged nuclear miniCic intensity (red) and FRET signal (purple) in the neighbours of dying cells. The curves were aligned on the peak of the FRET signal. Light areas are s.e.m.. Note that miniCic nuclear exclusion correlates well with the pulses of FRET activity (although there is a delay in the relocation of miniCic to the nucleus). **c:** Snapshots of local projections in the larval accessory cells (LECs) of the pupal abdomen expressing E-cad-GFP and miniCic. Blue stars show an extruding cell, blue arrows show the nuclei of neighbouring cells (which lose transiently miniCic signal). Dotted lines show cell contours. Scale bar=10μm. **d:** Averaged nuclear miniCic signal in the LECs neighbouring extruding cells (Time 0 is the termination of extrusion). Light areas are s.e.m.. **e:** Snapshots of a local projection of a pupal notum with *UAS-EGFR dsRNA* clones (blue, UAS-His3-mIFP) expressing E-cad-GFP and miniCic. White stars show two EGFR RNAi cells extruding. Blue lines show the contour and the nuclei of two EGFR RNAi cells neighbouring the dying cells, white lines show the contours and nuclei of WT cells neighbouring the dying cells. Note that dying EGFR depleted cells can still activate ERK in the WT neighbours (loss of nuclear miniCic) but ERK does not get activated in EGFR depleted cells. Scale bar=10μm.

**Figure S4:**
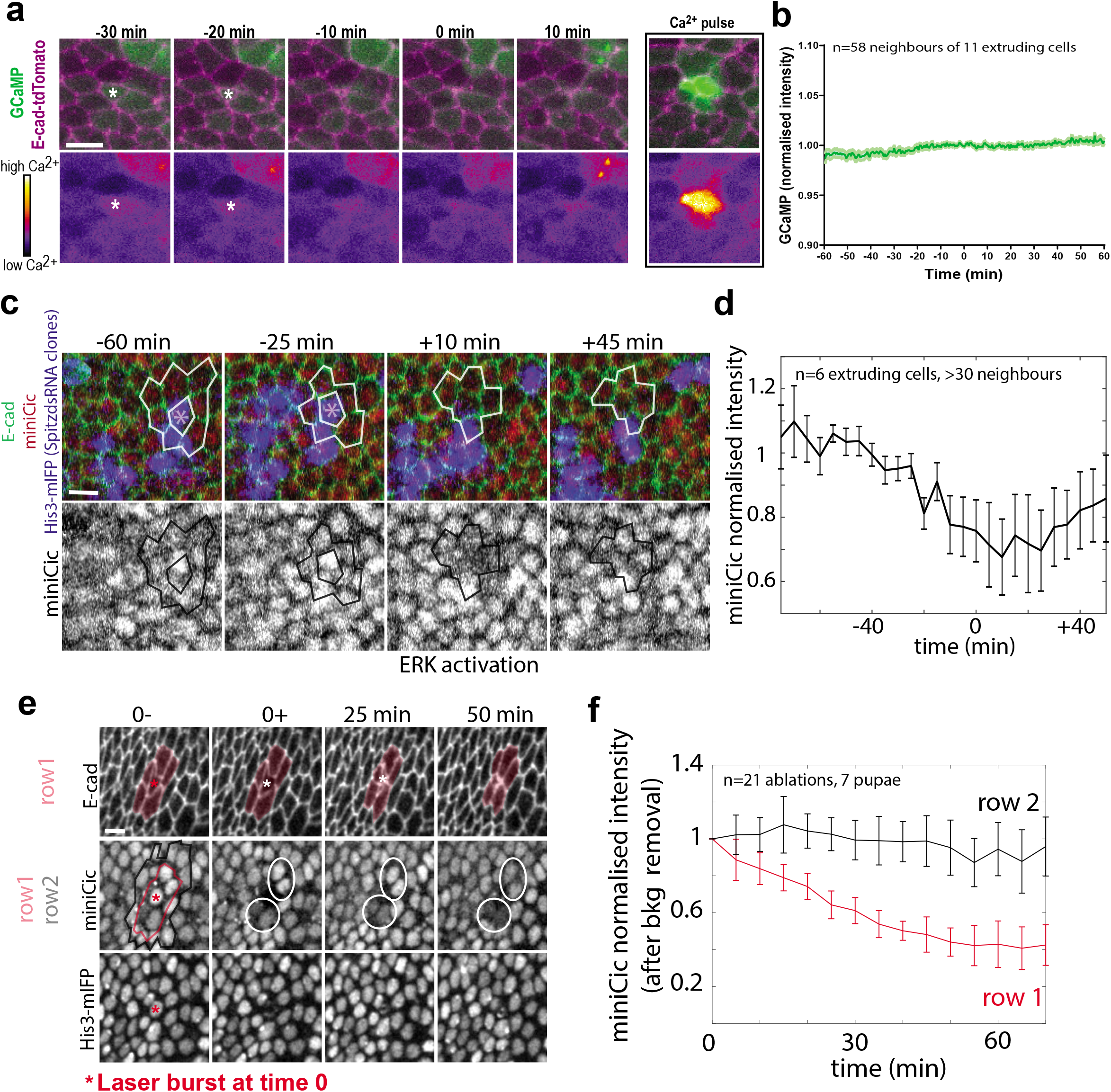
ERK pulses do not correlate with Ca^2+^ pulses, do not require Spitz expression in the dying cells, and can be mimicked by wound healing-induced stretching. **a:** Snapshots of local projections of a pupal notum expressing the Calcium sensor GCaMP (pseudo colour) and E-cad-tdTomato. The white stars show a dying cell. There is no visible change of Ca^2+^ activity in the neighbours or in the dying cell. The right inset show an example of rare spontaneous pulse of Ca^2+^ (one or two per movie, no correlation with cell death). Scale bar=10μm. **b:** Averaged and normalised GCaMP signal in cells neighbouring extruding cells (Time 0, termination of extrusion). Light area is s.e.m.. **c:** Snapshots of a local projection of a pupal notum with *UAS-spitz dsRNA* clones (blue, UAS-His3-mIFP) expressing E-cad-GFP and miniCic. The stars show a dying Spitz RNAi cell, white lines show the contour of the neighbours. Note that the Spitz RNAi effect was validated using the pnr-gal4 driver (nota fusion defects and reduction of ERK assessed with miniCic, not shown). Scale bar=10μm. **d:** Averaged and normalised miniCic nuclear intensity in cells neighbouring Spitz RNAi extruding cells (Time 0, termination of extrusion). Error bars are s.e.m.. **e:** Snapshots of local projection of a pupae expressing E-cad-GFP, miniCic and His3-mIFP. The cell marked with a red star is laser ablated at time 0. Pink line/area show the first neighbours, black line the second row of neighbours. The white circles show miniCic signal in the direct neighbours of the laser ablated cell. Scale bar=10μm. **f:** Averaged and normalised nuclear miniCic signal (after background removal, acquisition on a spinning disc) in the first row (red) and second row (black) of cells neighbouring the laser ablated cells. Time 0 is the ablation time.

**Figure S5:**
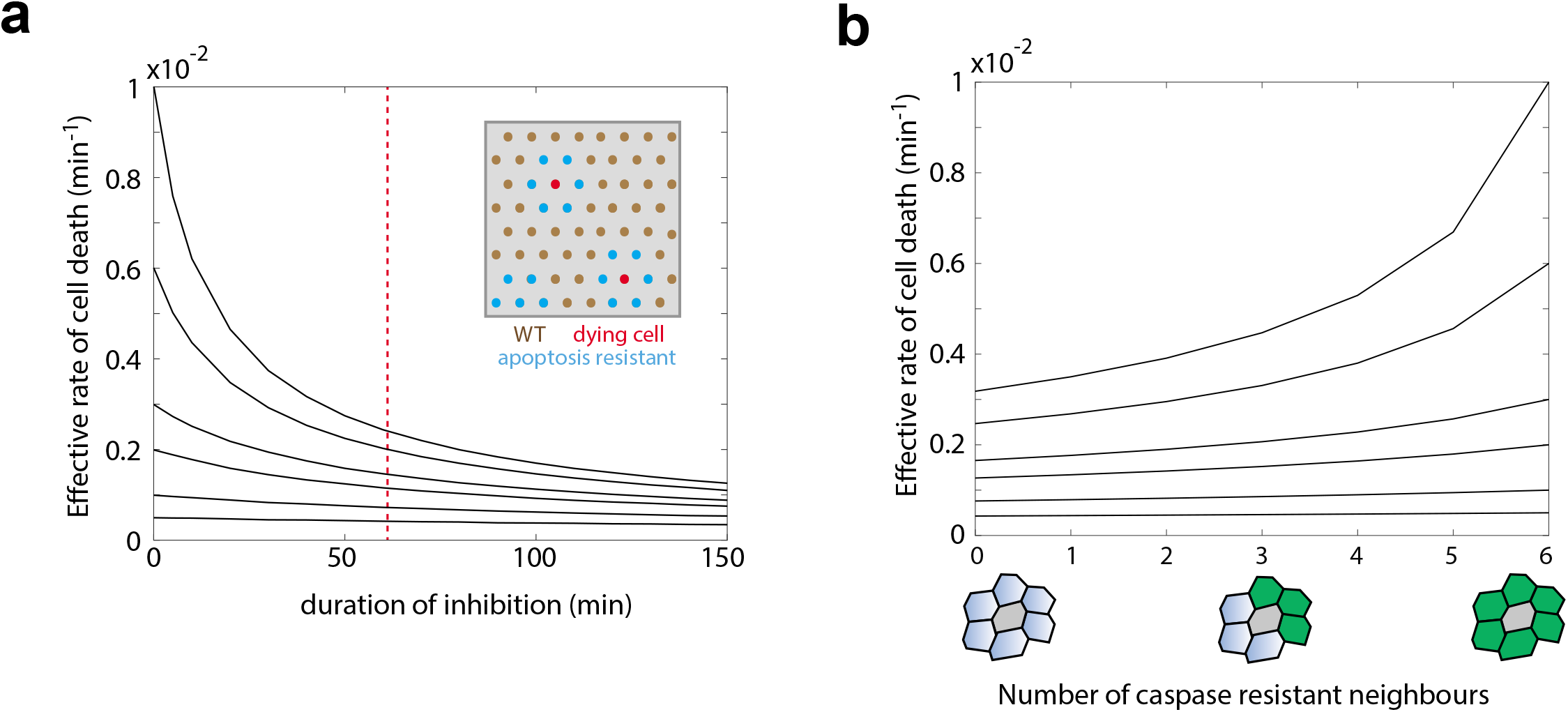
ERK pulses can modulate significantly the rate of cell death and could recapitulate features of cell competition. **a:** 2D simulations of the distribution of cell death assuming a local refractory phase of various times (see **Methods**) and different “intrinsic” rate of cell elimination (elimination rate without refractory phase, from bottom to top curves, λ=0.05, 0.1, 0.2, 0.3, 0.6, 1 x10^−2^ death.min^−1^). For a refractory phase of 60 minutes (red doted line) the curve showing a rate of cell elimination similar to the one measured in the posterior region of the pupal notum (~0.0013/minute) corresponds to an intrinsic rate of cell death λ=0.002. **b:** Estimation of the effective cell death rate in a cell (grey) neighbouring various number of caspase resistant cells (0 to 6, green cells) with different “intrinsic” rate of cell elimination (from bottom to top, λ=0.05, 0.1, 0.2, 0.3, 0.6, 1 x10^−2^ death.min^−1^). See **Methods**. In a region with a high rate of cell elimination, clones resistant for apoptosis will increase the rate of neighbouring cell elimination, especially for cells surrounded by several apoptosis resistant neighbours (similar to a cell competition scenario).

## Video legends

**Video S1: Induction of clone elimination by optoDronc is effector caspase-dependent**

Local projections of pupal nota expressing E-cad-tdTomato (green) and optoDronc (magenta, left) or optoDronc and p35 (magenta, right) in clones. Blue light exposure starts at the onset of the movies. Scale bars=10μm.

**Video S2: Induction of extrusion by optoDronc in different clone topologies**

Local projections of pupal nota expressing E-cad-tdTomato (green) and optoDronc (magenta) in clones of various sizes (one cell, two cells, three cells in raw, three cells in cluster, more than four cells in cluster). Blue light exposure starts at the onset of the movies. Scale bars=10μm.

**Video S3: optoDronc clone elimination and Dextran diffusion**

Local projections of pupal nota expressing E-cad-tdTomato (green, middle) and optoDronc (magenta) in clones upon injection of far red Dextran (grey levels, same plane as E-cad, right). Top part shows a line of four cells, bottom a cluster of four cells with aberrant extrusion. Blue light exposure starts at the onset of the movies. Scale bars=10μm.

**Video S4: Cell extrusion localisation in a WT notum**

Local projection of a pupal notum marked with E-cad-GFP showing every cell extrusion (coloured circles). Anterior: left, Posterior: right. Circles appear at the termination of extrusion (orange) and stay for nine frames (progressively becoming blue). Scale bar=10μm.

**Video S5: Reversion of caspase activity in a cell neighbouring an extruding cell**

Local projection of a pupal notum expressing GC3Ai (green and right, effector caspase activity) and E-cad-tdTomato (magenta). The bottom cell dies, while GC3Ai signal stop increasing in the top cell which remains in the tissue. Scale bar=10μm.

**Video S6: Transient activation of ERK in cells neighbouring extruding cells**

Local projection of a pupal notum expressing miniCic-mScarlet (magenta, bottom right, ERK sensor), E-cad-GFP (green, top right) and His3-mIFP (grey, bottom left). Cells marked in cyan are extruding. The white lines show the nuclei of the direct neighbouring cells (transient loss of nuclear miniCic= ERK activation). Scale bar=10μm.

**Video S7: Transient ERK activation is also visible with a FRET sensor**

Local projection of a pupal notum expressing miniCic-mScarlet (grey, left), and EKARnls (ERK FRET sensor, FRET ratio in pseudocolour, red=activation, blue=inhibition). The green circles show a dying cell. Note the transient loss of nuclear miniCic in the direct neighbours and the transient burst of FRET signal. Scale bar=10μm.

**Video S8: There is no visible Ca^2+^ pulses near extruding cells**

Local projection of a pupal notum expressing GCaMP (green, right), and E-cad-tdTomato (magenta). The cyan contours show an extruding cell. Scale bar=10μm.

**Video S9: Spitz is not required in the dying cell to activate ERK in the neighbours**

Local projection of a pupal notum expressing miniCic-mScarlet (magenta, bottom right, ERK sensor), E-cad-GFP (green, top right) and His3-mIFP in clones expressing Spitz-dsRNA (grey, bottom left). The cell in cyan is a Spitz RNAi cell dying. The white circles show the nuclei of direct neighbouring WT cells (transient loss of nuclear miniCic = ERK activation). Scale bar=10μm.

**Video S10: Activation of ERK driven by single cell laser ablation**

Local projection of a pupal notum expressing miniCic-mScarlet (magenta, bottom right, ERK sensor), E-cad-GFP (green, top right) and His3-mIFP (grey, bottom left). The cell in cyan is ablated by a UV pulsed laser at time 0. The white circles show the nuclei of direct neighbouring cells (transient loss of nuclear miniCic = ERK activation). Scale bar=10μm.

**Video S11: EGFR is required for extrusion-induced ERK activation**

Local projection of a pupal notum depleted for EGFR (*pnr-gal4, UAS-egfrdsRNA*) expressing miniCic-mScarlet (red, bottom left, ERK sensor), E-cad-GFP (green, top right) and His3-mIFP (blue, bottom right). White contours show the dying cell. The white circles show the nuclei of direct neighbouring cells (no change of nuclear miniCic, no ERK activation). Scale bar=10μm.

**Video S12: ERK is also activated by cell extrusion in the pupal abdomen**

Local projection of larval accessory cells in the pupal abdomen expressing miniCic-mScarlet (magenta, grey on the right, ERK sensor) and E-cad-GFP (green). The red contours show the dying cell. The blue contours show the direct neighbours. Scale bar=10μm.

**Video S13: ERK pulses precede effector caspase inhibition**

Local projection of a pupal notum expressing GC3Ai (green and right, effector caspase activity) and miniCic (magenta, middle). Green circles track the nucleus of the first dying cell. Purple circles track the nucleus of a neighbour undergoing transient caspase inhibition (slow-down of GC3Ai signal increase). Scale bar=10μm.

**Video S14: EGFR depletion abolishes caspase reversions**

Local projection of a pupal notum depleted for EGFR (*pnr-gal4, UAS-egfrdsRNA*) expressing GC3Ai (green and right, effector caspase activity) and E-cad-tdTomato (magenta). Coloured contours show neighbouring cells activating caspase. Scale bar=10μm.

**Video S15: Cell extrusion localisation in a EGFR RNAi notum**

Local projection of a pupal notum depleted for EGFR (*pnr-gal4, UAS-egfrdsRNA*) marked with E-cad-GFP showing every cell extrusion (coloured circles). Anterior: left, Posterior: right. Circles appear at the termination of extrusion (orange) and stay for nine frames (progressively becoming blue). Scale bar=10μm.

**Video S16: Clusters of cell elimination and aberrant extrusions appear upon EGFR depletion**

Local projection from different pupal nota depleted for EGFR (*pnr-gal4, UAS-egfrdsRNA*) marked with E-cad-GFP showing four examples of clusters of cell extrusion (cyan, more than four cells) followed by aberrant extrusions (relaxation, wound healing and E-cad accumulation at vertices). Scale bar=10μm.

